# Higher-order inter-chromosomal hubs shape 3-dimensional genome organization in the nucleus

**DOI:** 10.1101/219683

**Authors:** Sofia A. Quinodoz, Noah Ollikainen, Barbara Tabak, Ali Palla, Jan Marten Schmidt, Elizabeth Detmar, Mason Lai, Alexander Shishkin, Prashant Bhat, Vickie Trinh, Erik Aznauryan, Pamela Russell, Christine Cheng, Marko Jovanovic, Amy Chow, Patrick McDonel, Manuel Garber, Mitchell Guttman

**Affiliations:** Division of Biology and Biological Engineering, California Institute of Technology, Pasadena, CA 91125; Program in Bioinformatics and Integrative Biology, University of Massachusetts Medical School, Worcester, MA 01655; David Geffen School of Medicine, University of California, Los Angeles, Los Angeles, CA 90095, USA; Department of Biostatistics and Informatics, Colorado School of Public Health, Aurora, CO 80045; Department of Biology, Boston University, Boston, MA 02215; Department of Biological Sciences, Columbia University, New York, NY 10027; Program in Molecular Medicine, University of Massachusetts Medical School, Worcester, MA 01655

## Abstract

Eukaryotic genomes are packaged into a 3-dimensional structure in the nucleus of each cell. There are currently two distinct views of genome organization that are derived from different technologies. The first view, derived from genome-wide proximity ligation methods (e.g. Hi-C), suggests that genome organization is largely organized around chromosomes. The second view, derived from *in situ* imaging, suggests a central role for nuclear bodies. Yet, because microscopy and proximity-ligation methods measure different aspects of genome organization, these two views remain poorly reconciled and our overall understanding of how genomic DNA is organized within the nucleus remains incomplete. Here, we develop Split-Pool Recognition of Interactions by Tag Extension (SPRITE), which moves away from proximity-ligation and enables genome-wide detection of higher-order DNA interactions within the nucleus. Using SPRITE, we recapitulate known genome structures identified by Hi-C and show that the contact frequencies measured by SPRITE strongly correlate with the 3-dimensional distances measured by microscopy. In addition to known structures, SPRITE identifies two major hubs of inter-chromosomal interactions that are spatially arranged around the nucleolus and nuclear speckles, respectively. We find that the majority of genomic regions exhibit preferential spatial association relative to one of these nuclear bodies, with regions that are highly transcribed by RNA Polymerase II organizing around nuclear speckles and transcriptionally inactive and centromere-proximal regions organizing around the nucleolus. Together, our results reconcile the two distinct pictures of nuclear structure and demonstrate that nuclear bodies act as inter-chromosomal hubs that shape the overall 3-dimensional packaging of genomic DNA in the nucleus.

## INTRODUCTION

Although the same genomic DNA is packaged in the nucleus of each cell, different sets of genes are expressed in different cell states^1,2^. Despite significant progress over the past decade, there are still many unanswered questions about how the genome is organized within the nucleus and how this changes across different cell states^3–7^. For example, it remains unclear whether inter-chromosomal interactions play an important role in shaping genome organization^8–14^.

There are currently two distinct and poorly reconciled views of genome organization that are derived from different technologies. The first view is primarily derived from genome-wide proximity ligation methods, which work by ligating the ends of DNA regions that are in close spatial proximity in the nucleus followed by sequencing to map interactions (e.g. 3C, Hi-C, ChIA-PET)^15–19^. In this view, genome organization is largely organized around chromosome territories, such that most DNA interactions occur within an individual chromosome^10,20–22^. These interactions include chromatin loops that connect specific genomic DNA regions such as enhancers and promoters^23–25^, local interacting neighborhoods of DNA called topologically associated domains (TADs)^26–28^, and compartments where DNA regions interact based on their transcriptional activity (A/B compartments)^5,9,29,30^. However, the extent to which DNA interactions occur between chromosomes has been controversial^9,10,20,24,31,32^.

The second view is primarily derived from *in situ* imaging of DNA, RNA, and protein in the nucleus using microscopy. In this view, the genome is also organized around structures such as nuclear bodies that typically concentrate DNA, RNA, and protein molecules that are associated with shared functional or regulatory roles within the nucleus^33–37^. These include nuclear bodies associated with ribosomal RNA transcription, processing, and biogenesis (nucleolus)^16,33,38^, spliceosomal complex assembly (Cajal bodies)^34,39,40^, and storage of mRNA processing and splicing factors (nuclear speckles)^41–44^, among others^35,45,46^. There is evidence that specific inter-chromosomal interactions can occur at these nuclear bodies. For example, nucleoli are formed around the active transcription of ribosomal DNA genes that are present across several distinct chromosomes^33,38,47^. In addition, specific actively transcribed genes from different chromosomes can localize near the periphery of nuclear speckles^48–51^. These observations, and others^8,11,13,52–54^, demonstrate that genome interactions can occur beyond chromosome territories^10,20,48,55^. Yet, because microscopy is limited to the analysis of a small number of DNA sites simultaneously, it remains unclear whether these observations represent special cases involving a small number of genomic sites or broader principles of genome organization.

There is a growing appreciation that microscopy and proximity-ligation methods can fail to identify similar genome structures^56–61^ because these two methods measure different aspects of genome organization. Specifically, microscopy measures the 3-dimensional spatial distances between DNA sites within single cells, whereas proximity-ligation methods measure the relative frequency with which two DNA sites are close enough in the nucleus to directly ligate^59^. This discrepancy is particularly significant when considering DNA regions that are organized around nuclear bodies, which can range in size from 0.5-2 μm^35,62^, and therefore may be too far apart to ligate. These technical issues highlight why our overall understanding of how genomic DNA is organized within the nucleus remains incomplete.

Here, we develop a method called Split-Pool Recognition of Interactions by Tag Extension (SPRITE) that moves away from the proximity-ligation paradigm and enables genome-wide detection of multiple DNA interactions that occur simultaneously within the nucleus. Using SPRITE, we recapitulate known genome structures identified by Hi-C, including compartments, topologically associated domains, and loop structures, and identify that many of these occur within higher-order structures in the nucleus. Importantly, we show that the contact frequencies measured by SPRITE strongly correlate with the 3-dimensional distances measured by microscopy. In addition to known structures, SPRITE identifies two major hubs of inter-chromosomal interactions that are spatially arranged around the nucleolus and nuclear speckles, respectively. Interestingly, we find that the majority of genomic regions exhibit preferential spatial association relative to one of these nuclear bodies, with regions that are highly transcribed by RNA Polymerase II organizing around nuclear speckles and transcriptionally inactive and centromere-proximal regions organizing around the nucleolus. Together, our results reconcile these two distinct pictures of nuclear structure and demonstrate that nuclear bodies act as inter-chromosomal hubs that shape the overall 3-dimensional packaging of genomic DNA in the nucleus.

## RESULTS

### SPRITE: A genome-wide method to identify higher-order DNA interactions in the nucleus

We sought to develop a genome-wide method that measures 3-dimensional spatial distances between DNA sites and enables mapping of higher-order interactions that occur simultaneously between multiple DNA sites within the same nucleus. To do this, we developed SPRITE, a method that moves away from proximity ligation. SPRITE works as follows: DNA, RNA, and protein are crosslinked in cells, nuclei are isolated, chromatin is fragmented, interacting molecules within an individual complex are barcoded using a split-pool strategy, and interactions are identified by sequencing and matching all reads that contain identical barcodes (Figure 1A, **see Methods**).

**Figure 1.**
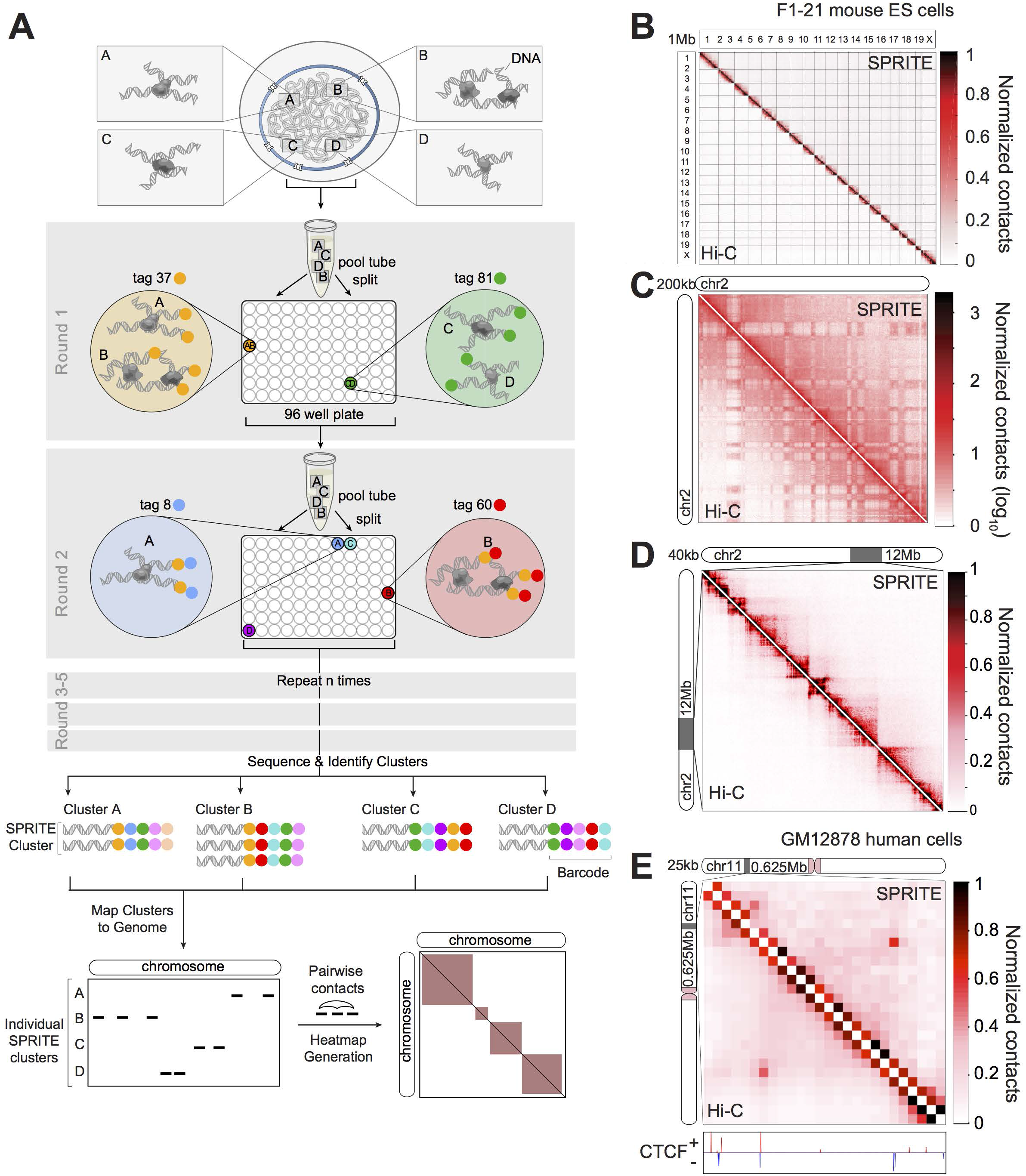
SPRITE accurately maps known genome structures across various resolutions. (A) A schematic of the SPRITE protocol. Crosslinked DNA complexes are split into a 96-well plate and molecules in each well are tagged with a unique sequence. After ligation, the molecules are pooled. This split-pool process is iterated and tags are sequentially added. The tagged DNA is sequenced and all reads with the same tag sequences (barcode) are matched into groups (SPRITE cluster). SPRITE clusters are converted to pairwise contacts or analyzed as individual clusters. (B-D) Pairwise contact maps comparing SPRITE (upper diagonal) and Hi-C (lower diagonal) in mouse embryonic stem (mES) cells at 1Mb resolution (B), on chromosome 2 at 200kb resolution (shown in log scale to improve visualization of long-range compartments in both Hi-C and SPRITE), (C) and in a 12Mb region on chromosome 2 at 40kb resolution (D). (E) Comparison of SPRITE and Hi-C contact maps in human GM12878 lymphoblast cells showing a chromatin loop at 25kb resolution. CTCF ChIP-seq peaks (ENCODE) are shown according to the positive (red) or negative (blue) orientation of the CTCF consensus motif.

Specifically, we uniquely barcode each molecule in a crosslinked complex by repeatedly splitting all complexes across a 96-well plate (“split”), ligating a specific tag sequence onto all DNA molecules within each well (“tag”), and then pooling these complexes into a single well (“pool”). After several rounds of split-pool tagging, each molecule in an interacting complex contains a unique series of ligated tags, which we refer to as a barcode (Figure 1A). Because all molecules in a crosslinked complex are covalently linked, they will sort together in the same wells throughout each round of the split-pool tagging process and will contain the same barcode, whereas the molecules in separate complexes will sort independently and therefore will obtain distinct barcodes. Therefore, the probability that molecules in two independent complexes will receive the same barcode decreases exponentially with each additional round of split-pool tagging (see **Methods**). For example, after 6 rounds of split-pool tagging, there are ∼10^12^ possible unique barcode sequences, which exceeds the number of unique DNA molecules present in the initial sample (∼10^9^). After split-and-pool tagging, we sequence all tagged DNA molecules and match all reads with shared barcodes. We refer to all unique DNA reads that contain the same barcode as a SPRITE cluster.

To confirm that DNA reads within a SPRITE cluster represent interactions that occur in the nucleus and are not formed by spurious association or aggregation in solution, we mixed crosslinked lysates from human and mouse cells prior to performing SPRITE and found that ∼99.8% of all SPRITE clusters contained human or mouse reads, but not both (see **Methods, Figure M1**). We ensured that the majority of DNA molecules within a crosslinked complex are barcoded by optimizing the ligation efficiency such that >90% of DNA molecules are ligated during each round of split-pool tagging (Figure S1A).

SPRITE differs from previous methods in several ways. In contrast to Hi-C, SPRITE can measure multiple DNA molecules that simultaneously interact within an individual nucleus, provides information about interactions that are heterogeneous from cell-to-cell, and measures contacts that are proportional to 3-dimensional spatial distance in the nucleus. In contrast to Genome Architecture Mapping (GAM)^63^, another proximity-ligation independent method, SPRITE can be performed without requiring specialized laboratory equipment or training, is significantly cheaper and faster to perform, and does not require extensive whole genome amplification. Furthermore, because SPRITE does not rely on proximity-ligation or whole genome amplification, it can be extended beyond DNA to directly incorporate RNA localization simultaneously.

### SPRITE accurately maps known genome structures across various resolutions

To test whether SPRITE can accurately map genome structure, we compared the results obtained by SPRITE to those measured by Hi-C^21^. Specifically, we generated SPRITE maps in two mammalian cell types that have been previously mapped by Hi-C – mouse embryonic stem cells (mES)^26^ and human lymphoblastoid cells (GM12878)^23^. We generated ∼1.5 billion sequencing reads from each sample and matched reads containing the same barcode to obtain ∼50 million SPRITE clusters from each sample (see **Methods**). These SPRITE clusters range in size from 2 reads to more than 1000 reads per cluster (Figure S1B). To directly compare SPRITE and Hi-C, we converted SPRITE clusters into pairwise contact frequencies by enumerating all pairwise contacts observed within a single cluster and down-weighting each pairwise contact by the total number of molecules contained within the cluster (Figure 1A, **Methods**). This normalization prevents the large SPRITE clusters from disproportionately impacting the pairwise frequency maps by ensuring that the number of contacts is linearly proportional to the number of reads in a cluster (see **Methods**).

Overall, the pairwise contact maps generated with SPRITE are highly comparable to Hi-C maps, with similar structural features observed across all levels of genomic resolution. At a genome-wide level, we observe a clear preference for interactions that occur within the same chromosome (Figure 1B). At a chromosome-wide level, we observe an alternating interaction pattern that is highly correlated with A and B compartments identified by Hi-C in both the human and mouse data (Spearman ρ= 0.85, 0.93, respectively) (Figure S1C-E). These regions have been shown to correspond to locations of active and inactive transcription^24,64^. At 40kb resolution, we observe that compartments are divided into topologically associated domains (TADs), where adjacent DNA sites organize into highly self-interacting domains that are separated by boundaries that preclude interactions with other neighboring regions (Figure 1D). We find that the location and strength of TAD boundaries, measured by the insulation scores across the genome, are highly correlated in Hi-C and SPRITE for both human and mouse (Spearman ρ= 0.90, 0.94) (Figure S1F-H). Finally, at 25kb resolution, we observe specific “looping” interactions that connect local regions that contain the expected convergent CTCF motif orientation previously described for loop structures (Figure 1E, Figure S1I)^23,65,66^. More generally, we find that the loops previously identified by Hi-C are strongly enriched within our SPRITE data at 10kb resolution (Figure S1I-K, see **Methods**).

Together, these results demonstrate that SPRITE generates accurate maps of genome structure across multiple levels of resolution.

### SPRITE identifies higher-order interactions that occur simultaneously

In addition to confirming pairwise genome structures identified by Hi-C, SPRITE can also directly measure multiple DNA regions that interact simultaneously within an individual cell, which we refer to as higher-order interactions. Although other proximity-ligation methods have been recently reported that can also map higher-order interactions^67,68^, these approaches are largely restricted to mapping 3-way or 4-way contacts. In contrast, SPRITE is not restricted in the number of higher-order contacts it can identify.

To explore the higher-order structures identified by SPRITE, we enumerated all interactions that occur simultaneously between 3 or more independent genomic regions (*k* ≥ 3), which we refer to as a *k*-mer. We found that one of the largest determinants of *k*-mer frequency in our data was the linear genomic distance separating each region in the *k*-mer. To account for this, we computed an enrichment score by normalizing the frequency of the observed *k-*mer by the average frequency observed across random *k*-mers that retain the same genomic distance (Figure S2A). We excluded *k-*mers that contain genomic regions that significantly deviate in coverage level relative to the average genomic level (see **Methods**).

Overall, we identified >310,000 *k-*mers (1Mb resolution, *k*=3-14 regions) that were observed in at least 5 independent SPRITE clusters, occurred at a frequency that exceeded 90% of the random permutations, and occurred >4-fold more frequently than the average of the permuted regions (Figure S2B, **Table S1,** See **Methods**). These enriched *k*-mers include various higher-order genomic DNA structures, including active compartments, gene clusters, and consecutive loop structures.

**(i) Active Compartments.** We observed highly enriched *k*-mers that connect multiple A compartment (transcriptionally active) regions that are non-contiguous and span large distances of the same chromosome. Specifically, we observed tens of thousands of SPRITE clusters that contain reads from at least three different A compartment regions that span at least 100Mb within an individual chromosome (max enrichment = 5.34, percentile = 100%, **Table S2**, Figure 2A, S2C).

**Figure 2.**
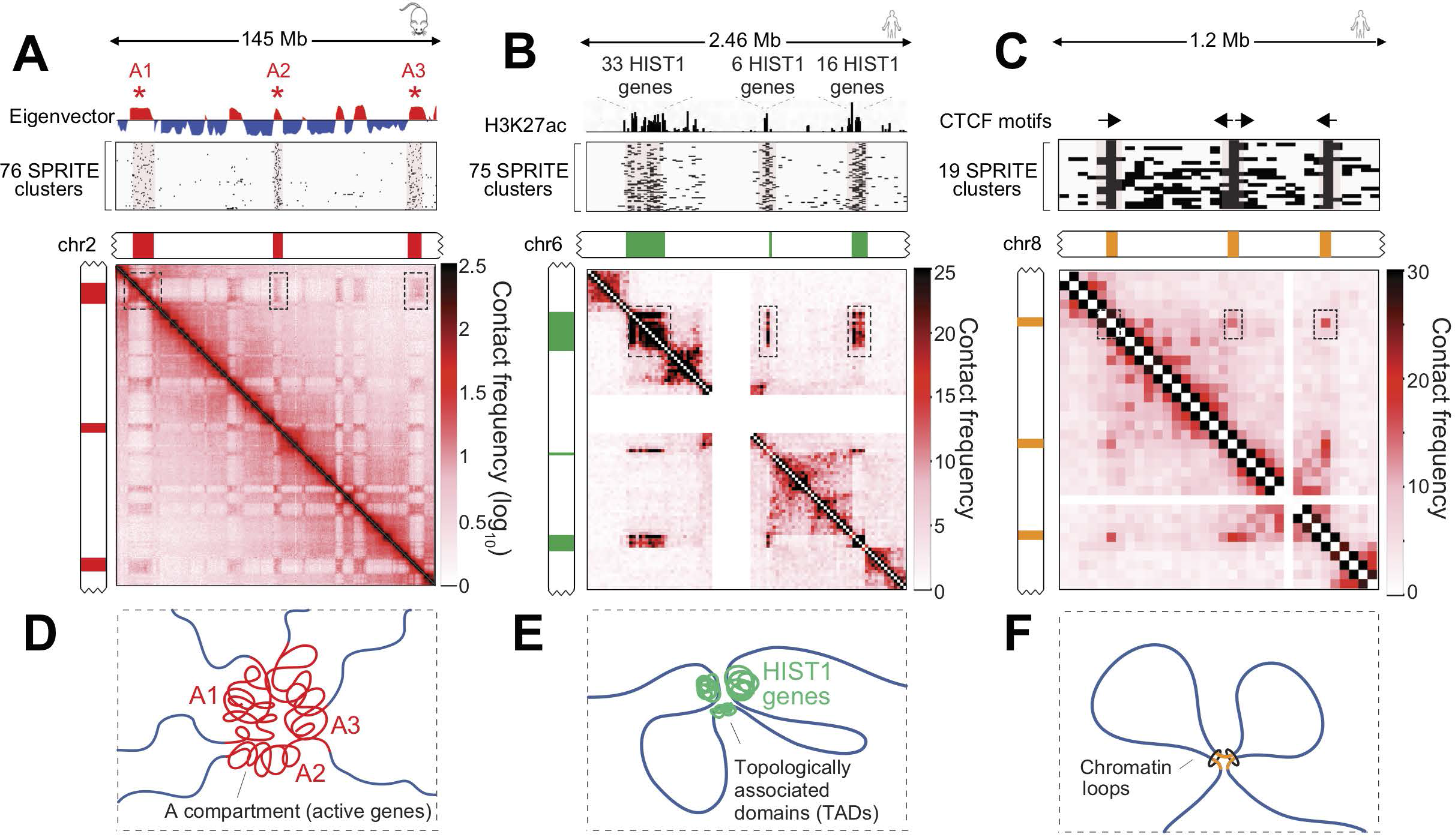
SPRITE measures higher-order interactions that occur simultaneously within the same nucleus. (A) Higher-order interactions between A compartment regions on chromosome 2 in mESCs. (Top) Compartment eigenvector showing A (red) and B (blue) compartments on chromosome 2. (Middle) Individual SPRITE clusters containing reads mapping to at least 3 distinct A compartment regions spanning over 100Mb binned at 1Mb resolution. Rows show individual SPRITE clusters and black lines denote 1Mb bins with at least 1 read within these clusters. (Bottom) SPRITE pairwise contact map of chromosome 2 at 200kb resolution. (B) Higher-order interactions between 3 TADs containing 55 histone genes in human GM12878 cells. (Top) H3K27ac ChIP-seq signal across a 2.46 Mb region on human chromosome 6. (Middle) 75 individual SPRITE clusters containing reads in all 3 TADs binned at 25kb resolution. (Bottom) SPRITE contact map of chromosome 6 from 25.70 to 28.16 Mb at 25kb resolution. Low coverage bins are masked. (C) Higher-order interactions between 3 loop anchors in human GM12878 cells. (Top) CTCF motifs and orientations at the 3 loop anchors. (Middle) 19 SPRITE clusters containing reads at all three loop anchors binned at 25kb resolution. (Bottom) SPRITE contact map of human chromosome 8 from 133.5 to 134.8 Mb at 25kb resolution. (D-F) Cartoon representations of higher-order interactions observed in SPRITE clusters between (D) A compartment regions, (E) HIST1 gene clusters and (F) consecutive loop anchors.

**(ii) Gene Clusters.** We identified >75 SPRITE clusters that connect multiple non-contiguous genes that are contained within the human histone gene clusters (enrichment = 8.4-fold, percentile = 100%, Figure 2B). Specifically, the *HIST1* gene clusters span ∼2 Mb and contains three distinct clusters of genes that are interspersed by non-histone gene containing sequences^69,70^. Notably, these clusters skip the intervening transcriptionally inactive regions. The frequency of SPRITE clusters that connect the three histone gene clusters was never observed in any of the 100 randomly permuted *k*-mers containing the same genomic distance.

**(iii) Consecutive Loops**. We identified higher resolution structures that correspond to simultaneous interactions between consecutive loops. Specifically, previous Hi-C studies suggested that consecutive loops may form higher-order interactions that bring together three distinct regions of the genome^71^. Consistent with this, we observe several examples of highly enriched *k*-mers that correspond to consecutive loop structures (Figure S2D). For example, we observe >19 SPRITE clusters that contain reads corresponding to three loop anchor points on human chromosome 8 (enrichment = 10.1-fold, percentile=100%, Figure 2C).

Taken together, these results indicate that SPRITE can detect multiple DNA interactions that occur simultaneously within single cells. These include the observations that active DNA regions interact within higher-order compartments (Figure 2D), clusters of genes with shared function can associate with each other across non-contiguous genomic distances (Figure 2E), and that multiple consecutive loops can occur simultaneously within the same cell (Figure 2F).

### SPRITE identifies interactions that occur across large genomic distances

In analyzing the higher-order *k*-mers, we noticed that many highly-enriched examples correspond to intra-chromosomal and inter-chromosomal interactions that occur across large genomic distances. Because we did not observe these interactions in our normalized pairwise contact maps (Figure 1D, S3B), we hypothesized that SPRITE clusters of different sizes might provide distinct information about genome structure.

To explore this, we stratified the SPRITE clusters based on their number of reads and generated pairwise contact maps. We noticed that the number of pairwise contacts observed between two genomic regions as a function of their linear genomic distance decreases at strikingly different rates for the various cluster sizes (“distance decay rate”, Figure 3A). For small SPRITE clusters (2-10 reads), we observe a distance decay rate that is comparable to the rate observed by Hi-C, with most contacts occurring within close linear distances. Indeed, SPRITE clusters containing 2-10 reads also show pairwise contact maps that are highly comparable to Hi-C (Figure S3A). In contrast, for the larger SPRITE clusters (10-1000+ reads), we observed a larger number of contacts at longer genomic distances that corresponds to increasing cluster sizes (Figure 3A). These long-range interactions correspond to an increased number of contacts between specific genomic regions that are expected to interact. For example, we observe a significant increase in the number of interactions that occur between active compartments that are >100Mb apart (Figure 3B). These results demonstrate that the different SPRITE cluster sizes represent interactions that occur across various spatial distances in the nucleus.

**Figure 3.**
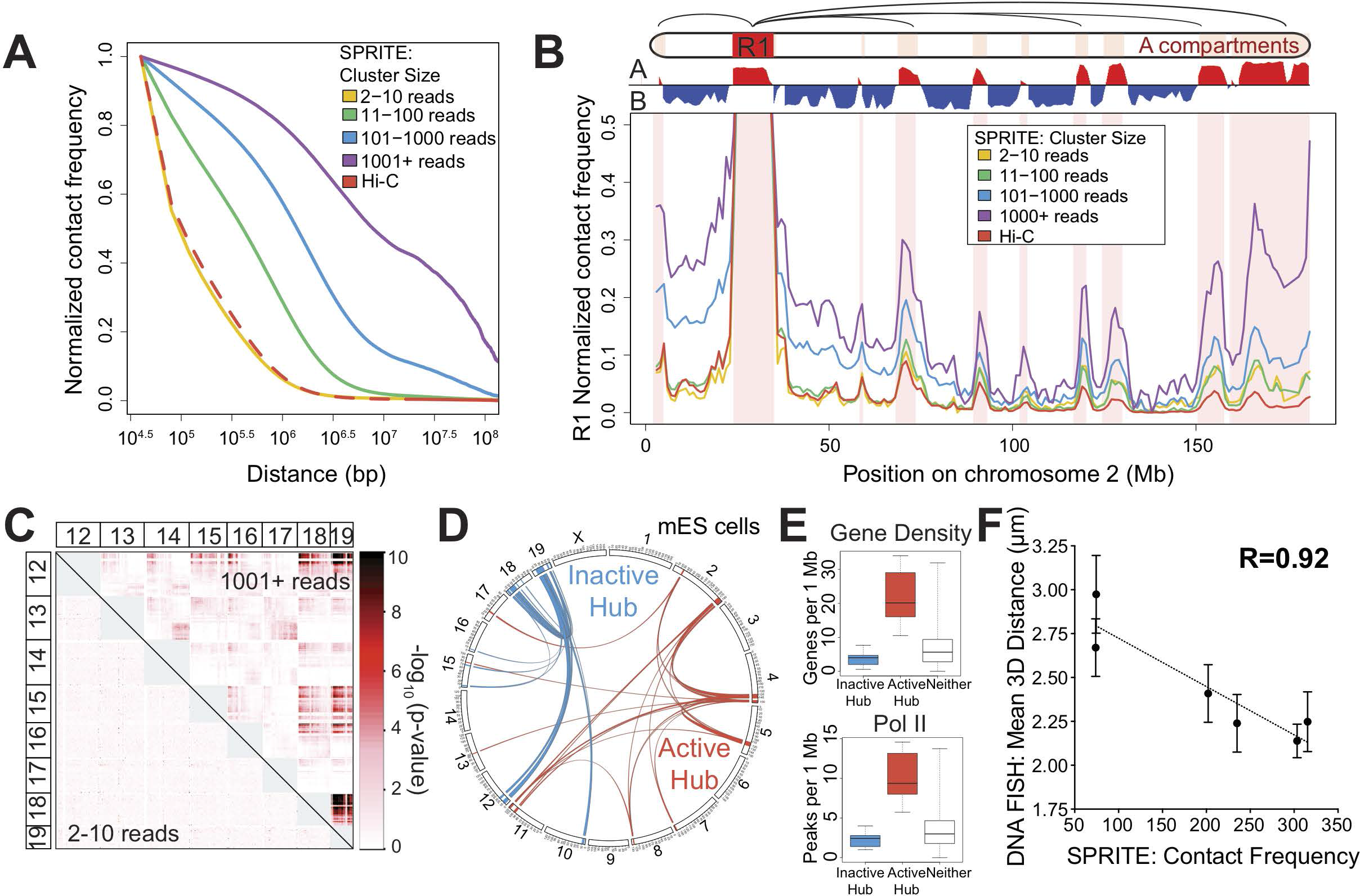
SPRITE measures interactions across large genomic distances and across chromosomes. (A) Relationship between contact frequency and linear genomic distance in mESCs. Contact frequencies are shown based on SPRITE clusters containing 2-10 reads (yellow), 11-100 reads (green), 101-1000 reads (blue) and 1001 or more reads (purple) and compared against Hi-C (red). (B) Contact frequency between a specific A (red) compartment region (R1: 25-34 Mb) and all other regions on mouse chromosome 2. Contact frequencies are shown for different SPRITE cluster sizes and compared against Hi-C. A and B compartment regions are shown in red and blue, respectively. (C) Inter-chromosomal contacts between chromosomes 12 through 19 in mES cells. Interaction *p*-values are shown for SPRITE clusters of size 2-10 reads (lower diagonal) and 1001+ reads (upper diagonal). *p*-values were calculated based on inter-chromosomal interaction frequencies and are shown in units of −log_10_(*p*-value). (D) Circos diagram of significant inter-chromosomal interactions in mES cells. Interactions are identified from unweighted inter-chromosomal contact maps in SPRITE clusters containing 2 to 1000 reads. Interactions with *p*-values less than 10^-10^ are shown. Two distinct networks of noninteracting connections are shown in blue (inactive hub) and red (active hub). (E) Box plots showing the distribution of gene density (top) and RNA polymerase II (Pol II) occupancy (bottom) for regions in the inactive hub, nucleolar hub, or neither hub. Gene density is calculated as genes per 1Mb and Pol II occupancy is calculated as number of ChIP-seq peaks per 1Mb. (F) Correlation between SPRITE inter-chromosomal contact frequency and 3D distance measured between six pairs of DNA sites (chr2A-chr11A; chr3A-chr14I; chr3A-chr19I; chr12I-chr15I; chr18I-chr19I; chr12I-chr19I) by 2-color DNA FISH.

### Inter-chromosomal interactions are partitioned into two distinct hubs

We also observed many inter-chromosomal interactions that occur at a greater frequency within the larger SPRITE clusters (10-1000+ reads) (Figure 3C, S3C-D). To explore these inter-chromosomal interactions, we built a graph connecting all 1Mb regions in the mouse genome containing a significant pairwise interaction (*p*-value < 10^−10^) (Figure 3D). These interactions segregate into two discrete “hubs” that contain distinct sets of mouse chromosomes and different functional activity. The first hub corresponds to gene poor and therefore transcriptionally inactive regions (“inactive hub”) and the second hub corresponds to gene dense regions that are highly transcribed (“active hub”, Figure 3E, S3E-F). Importantly, we observed two similar inter-chromosomal hubs in the human genome and these hubs displayed comparable functional activity as the hubs identified in the mouse genome (Figure S3G-J). Given the similar properties of the mouse and human inter-chromosomal hubs, we focused on mouse ES cells for our subsequent characterization of these hubs.

To test whether SPRITE accurately measures 3-dimensional spatial distances across chromosomes, we performed DNA FISH with probes against genomic DNA regions on 7 distinct chromosomes (see **Methods**). Importantly, we identified a striking correlation between the frequency measured by SPRITE and the average 3D distance measured between pairs of DNA regions by microscopy (Pearson correlation=0.922, Figure 3F). This demonstrates that SPRITE quantitatively measures the 3-dimensional spatial distance at which DNA sites on different chromosomes interact within the nucleus.

### The inactive inter-chromosomal hub is organized around the nucleolus

To understand where in the nucleus these inter-chromosomal interactions occur, we first explored the inactive hub and noticed that several of the genomic DNA regions in this hub are linearly close to regions that have been reported to contain ribosomal DNA^72–78^ (the regions in this hub do not contain rDNA themselves, see **Methods**). Because ribosomal DNA regions are known to be organized, transcribed, and processed within the nucleolus, we hypothesized that the genomic DNA regions in this hub may be organized around the nucleolus.

To test this, we explored whether the genomic DNA regions in this hub are associated with ribosomal RNA localization, which demarcates the nucleolus. Specifically, we adapted the SPRITE protocol to enable simultaneous mapping of interactions between RNA and DNA molecules (see **Methods**, Figure S4A). Using this approach, we mapped the interactions of ribosomal RNA on genomic DNA in mouse ES cells and found that it was specifically enriched over the genomic DNA regions contained within the inactive hub (Figure 4A). In fact, we found that ribosomal RNA enrichment across the genome is strongly correlated with how frequently a region is within the inactive hub (Figure 4A, S4B).

**Figure 4.**
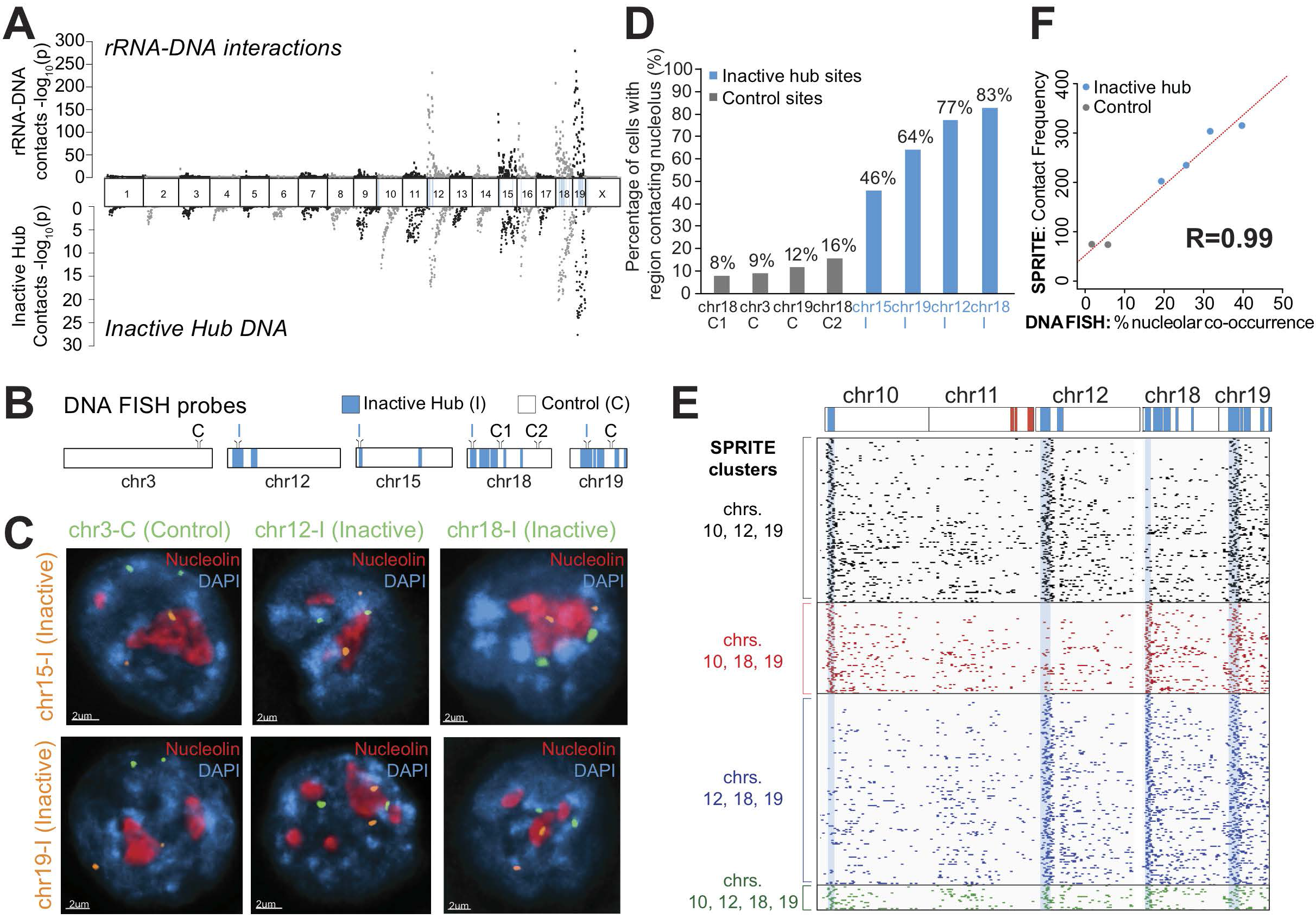
Genomic DNA in the inactive hub is organized around the nucleolus. (A) Ribosomal RNA (rRNA) localization across the mouse genome (top) compared to average inter-chromosomal contact frequency with regions in the inactive hub (bottom). *p*-values were calculated for rRNA localization and inactive hub contact frequency and are shown in units of – log_10_(*p*-value). DNA regions contained within the inactive hub are highlighted in blue in the chromosome ideogram. (B) Locations of probe regions used for DNA FISH experiments. Regions within the inactive hub are shown in blue. I denotes probe regions within the inactive hub and C denotes control probe regions outside the inactive hub. (C) Example images from immunofluorescence for nucleolin (red) combined with DNA FISH for six different pairs of DNA FISH probes (orange and green). Nucleolin was used to demarcate nucleoli and DAPI (blue) was used to demarcate nuclear DNA. (D) Comparison of DNA regions that directly contact the nucleolus for 8 different probe regions, including 4 control regions (grey) and 4 inactive hub regions (blue). Contact with the nucleolus for each probe region is calculated as the percent of cells with at least 1 allele that overlaps the nucleolin signal (distance = 0 μm) (n = 50 to 100 cells). (E) Example SPRITE clusters containing reads from different combinations of inactive hub regions on chromosomes 10, 12, 18 and 19 binned at 1Mb resolution. The ideogram at the top shows inactive hub regions in blue and active hub regions in red. Rows correspond to individual SPRITE clusters and dots denote 1Mb bins with at least 1 read. Three examples of 3-way interactions are shown in black, red and blue and a 4-way interaction is shown in green. (F) Comparison of SPRITE inter-chromosomal contact frequency and DNA FISH measurements of co-localization around the same nucleolus for six pairs of regions (see details in Figure S4C). The SPRITE contact frequency between two regions is shown (y-axis) relative to the percent of cells where at least one allele of both DNA regions co-localize at the same nucleolus as measured by DNA FISH (x-axis). Pairs of inactive hub regions are colored blue and pairs with one inactive hub region and one control region are colored grey.

To confirm that the inactive hub represents DNA sites physically located near the nucleolus *in situ*, we performed 3-dimensional DNA FISH combined with immunofluorescence for nucleolin, a well-known protein marker of the nucleolus. Specifically, we selected DNA FISH probes for 4 genomic regions that were identified in the inactive hub and 3 control regions that are on the same chromosomes but were not in the inactive hub. We selected an additional control region on a chromosome lacking any regions in the inactive hub (Figure 4B). We calculated the 3D distance between each allele and the nearest nucleolus and found that regions in the inactive hub are dramatically closer to the nucleolus than negative control regions (on average ∼750nm closer, Figure S4C). In the majority of the cells analyzed, at least one allele of the DNA regions in the inactive hub directly contacts the periphery of the nucleolus (∼61% of cells, Figure 4C-D, Figure S4C-D). Therefore, we refer to the DNA regions in this hub as the nucleolar hub.

Because there are many genomic regions in the nucleolar hub, we hypothesized that multiple DNA sites simultaneously interact around a single nucleolus. Consistent with this, we observed >1,200 SPRITE clusters that contain simultaneous interactions between at least three distinct genomic regions on different chromosomes in the nucleolar hub (max *k*-mer enrichment = 7.84-fold, median percentile =100%, Figure 4E, **Table S3**). To confirm that these inter-chromosomal contacts occur through co-localization at the same nucleolus, we performed 2-color DNA FISH combined with immunofluorescence for nucleolin and measured the frequency of co-association at the nucleolus (**Movie S1**). We observed that two regions in the nucleolar hub were >7-times more likely to co-occur around the same nucleolus compared to a nucleolar hub region and control region (Figure S4E). Importantly, the frequency of co-occurrence of DNA sites at the same nucleolus measured by DNA FISH is highly correlated with the frequency at which these genomic DNA regions co-occur in the SPRITE clusters (Pearson r=0.99, Figure 4F). This demonstrates that SPRITE quantitatively measures the frequency at which DNA sites co-occur in spatial proximity within single cells.

### The active inter-chromosomal hub is organized around nuclear speckles

We noticed that the genomic DNA regions within the active inter-chromosomal hub are strongly enriched for U1 spliceosomal RNA and Malat1 lncRNA localization (Figure S5A) and their localization levels are highly correlated with how frequently a DNA region interacts with the active hub (Spearman ρ =0.80 and 0.74, Figure 5A, S5B). Because U1 and Malat1 are known to localize at nuclear speckles^41,79–82^, a nuclear body that contains proteins involved in mRNA splicing and processing^83^, we hypothesized that inter-chromosomal interactions occurring between regions in the active hub may be spatially organized around nuclear speckles.

**Figure 5.**
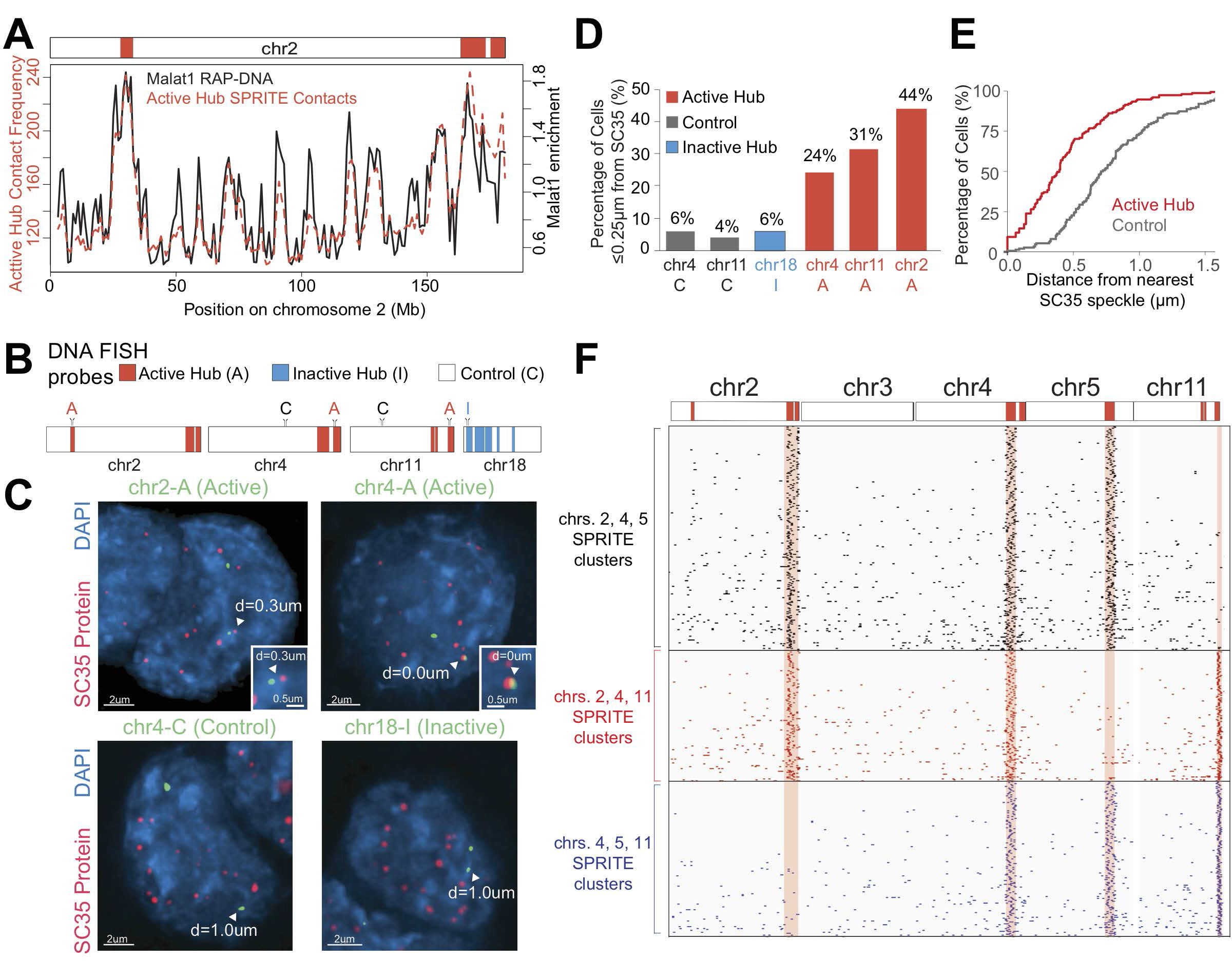
Genomic DNA in the active hub is organized around nuclear speckles. (A) Malat1 lncRNA localization across mouse chromosome 2 (black) compared to the average SPRITE inter-chromosomal contact frequency with regions in the active hub (red). Red boxes denote genomic regions within the active hub. (B) Locations of probe regions used for DNA FISH experiments. Regions within the active hub (“A”, red), regions in the inactive hub (“I”, blue), and control regions not present in either hub are shown in white (“C”). (C) Example images from immunofluorescence for SC35 (red) combined with DNA FISH for four DNA regions (green). SC35 foci were used to demarcate nuclear speckles (red) and DAPI was used to demarcate nuclear DNA (blue). Arrowheads denote one allele of the DNA FISH probe with the shortest 3D distance to SC35 and inset images show the region around the arrowhead. (D) Percentage of cells with at least 1 allele within 0.25 μm of an SC35 speckle (n = 50 to 100 cells). See further quantitation for all 6 DNA FISH probe distances in Figure S5D. (E) Cumulative frequency of the minimum 3D distances measured between all active hub (red) and control (grey) DNA FISH regions from their most proximal speckle, demarcated by SC35. See Figure S5E for CDF of all individual DNA FISH regions. (F) Example SPRITE clusters containing reads from different combinations of 3 active hub regions on chromosomes 2, 4, 5 and 11 binned at 1Mb resolution. The ideogram at the top shows active hub regions in red. Rows correspond to individual SPRITE clusters and black lines denote 1Mb bins with at least 1 read. Three different 3-way interactions are shown in black, red and blue.

To test this, we performed 3D DNA FISH combined with immunofluorescence for SC35, a well-known protein marker of nuclear speckles. We selected FISH probes targeting 3 DNA regions contained in the active hub and 2 control regions on the same chromosome that are not in the active hub. We also selected another control genomic region within the inactive hub (Figure 5B). We calculated the 3D distance between each region and the closest nuclear speckle and found that all 3 active hub regions are consistently closer than the 3 control regions to nuclear speckles (Figure 5C-E, S5C-E). Indeed, for genomic regions in the active hub, we observe a dramatic increase in the number of cells where at least one allele directly touches a nuclear speckle relative to control regions (∼13-fold, Figure S5D). Based on these observations, we refer to the DNA regions in this hub as the nuclear speckle hub.

We hypothesized that regions in the nuclear speckle hub may simultaneously associate with the same nuclear speckle. Indeed, we identified >690 SPRITE clusters containing at least three distinct active hub regions that were present on different chromosomes (max *k*-mer enrichment = 7.95-fold, median percentile =100%, **Table S4**, Figure 5F). Consistent with this, we observe that two DNA sites defined to be in the speckle hub by SPRITE are >8-times as likely to be within 1μm of each other compared to an active and control region by microscopy (Figure S5F-H). Notably, the number of SPRITE clusters containing multiple speckle hub regions is lower than for nucleolar hub regions, which may reflect the fact that nuclear speckles are both smaller in volume and present in larger numbers (>15/nucleus) within each nucleus compared to the nucleolus (∼1-3/nucleus).

These results demonstrate that actively transcribed genes that are present on multiple different chromosomes can form higher-order interactions that are spatially organized around nuclear speckles.

### Nuclear bodies shape the overall 3D organization of chromosomes in the nucleus

To understand how the remainder of the genome is organized in the nucleus relative to these nuclear bodies, we assessed whether genomic DNA regions that are not directly within the nucleolar or nuclear speckle hubs also show preferential 3-dimensional associations relative to these nuclear bodies. To test this, we calculated the average number of SPRITE contacts with nucleolar hub regions and the nuclear speckle hub regions for each 1Mb region in the genome (Figure 6A). We find that the majority of genomic regions exhibit preferential spatial association with either the nucleolus or nuclear speckles (Figure 6A, S6C). Interestingly, genomic DNA preferences for the nucleolus or nuclear speckles are mutually exclusive, such that genomic regions that interact frequently with the nucleolus are depleted relative to the nuclear speckles and vice versa (Figure 6A, S6B-C). We confirmed that these SPRITE contact frequencies accurately represent the 3-dimensional spatial distance relative to these nuclear bodies and found that they are very highly correlated with distances measured by microscopy (Pearson r = 0.98, 0.96) (Figure 6B, S6A).

**Figure 6.**
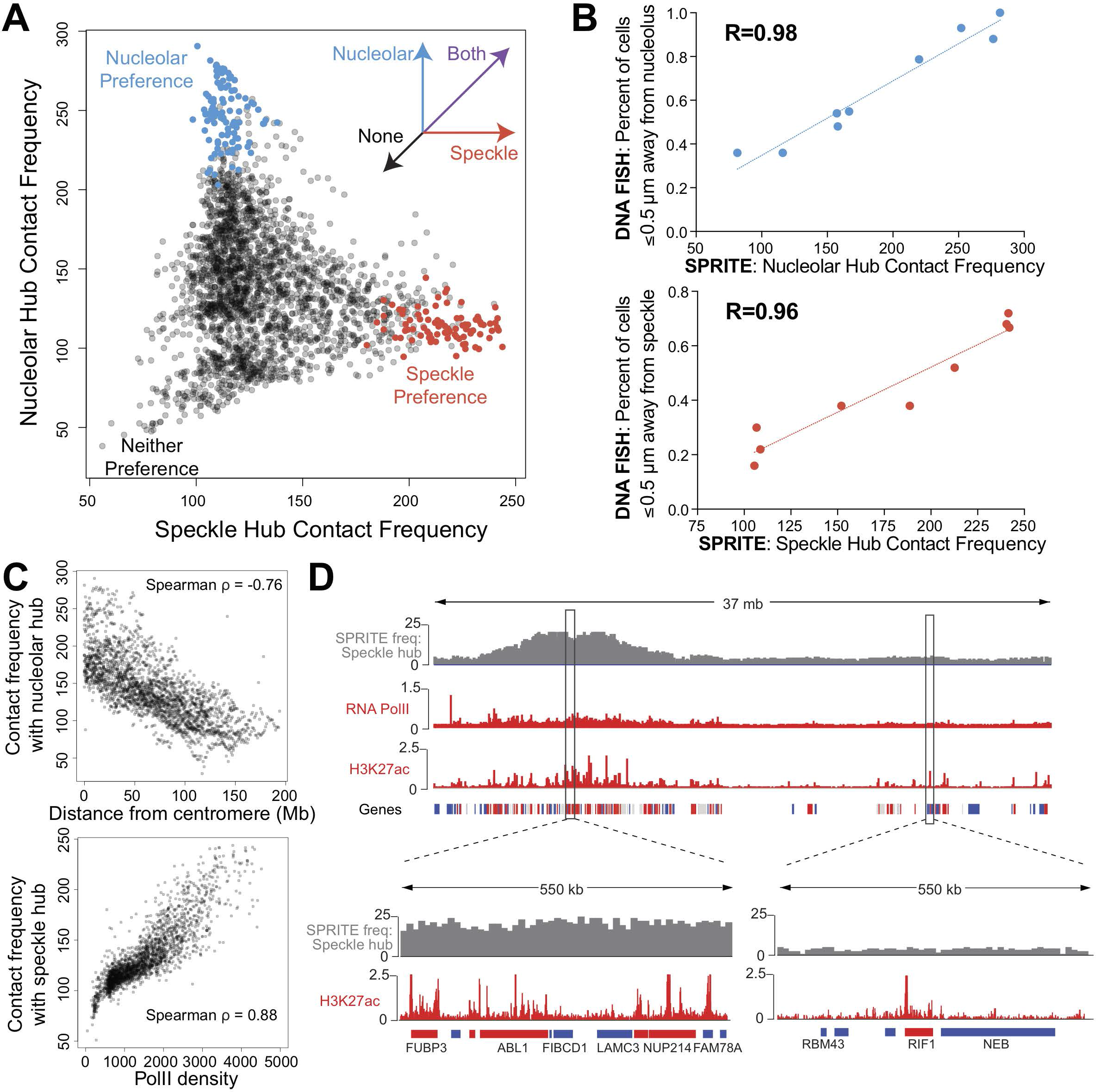
Preferential organization of DNA relative to the nucleolus and nuclear speckles shapes the overall organization of genomic DNA in the nucleus. (A) Genome-wide comparison of average SPRITE inter-chromosomal contact frequency to regions in the nucleolar hub (y-axis) or speckle hub (x-axis). Each dot corresponds to a 1Mb bin and colored dots denote DNA regions within the nucleolar hub (red) or speckle hub (blue). (B) SPRITE contact frequencies with the nucleolar hub (top) or speckle hub (bottom) regions (x-axis) compared to the DNA FISH contact frequency to nucleoli or nuclear speckles as measured by microscopy (y-axis). Microscopy distance to nuclear speckles and nucleoli are based on measurements between the closest allele per cell to the most proximal nuclear body using DNA FISH and IF experiments using SC35 and nucleolin as markers for nuclear speckles and nucleoli, respectively. (C) Comparison of the average SPRITE inter-chromosomal contact frequencies with the nucleolar hub and linear distance from the centromere (top) and with the speckle hub and PolII density (from ENCODE) (bottom). (D) Average speckle hub inter-chromosomal contact frequencies computed on chromosome 2 binned at 10kb resolution compared to RNA PolII and H3K27ac signal (ENCODE) for active transcription. Genes are demarcated as highly expressed (red), moderately expressed (grey), or inactive (blue) based on FPKM values (>10 high, 2-10 moderate, and 0-2 inactive). Speckle hub contact frequencies are zoomed in for actively transcribed gene dense region on chr2:31.4-30.0Mb and a gene poor transcriptionally inactive neighborhood on chr2:51.7-52.3Mb.

To understand the basis of these spatial preferences, we examined whether the genomic regions that are closer to a specific nuclear body correspond with various structural or functional properties.

**(i) Nucleolar preference.** We found that regions that are linearly close to the centromere are closer to the nucleolus (Figure 6C, Spearman ρ = 0.76). Notably, these results are consistent with previous microscopy and genomic observations that have shown that centromeres often co-localize on the periphery of the nucleolus^47,84–86^ (referred to as “chromocenters”). However, not all genomic regions that are close to centromeres are close to the nucleolus because actively transcribed genes are excluded from the nucleolar compartment even when they reside in linear proximity to a centromere (Figure S6E). These results indicate that genomic DNA that is organized around the nucleolus is largely gene poor and inactive chromatin, consistent with previous observations that the nucleolus can act as an anchor for inactive chromatin^87–89^. We do not observe a global relationship between regions that are in the nucleolar hub and regions that are associated with the nuclear lamina (Figure S7A, spearman ρ = 0.01), a distinct nuclear compartment that is also enriched for inactive chromatin, suggesting a precise organization pattern such that different inactive DNA regions associate preferentially with distinct nuclear compartments. While other studies have previously mapped individual regions that interact at the nucleolus^38,90^, our results provide the first 3-dimensional picture of how these DNA sites arrange around the nucleolus relative to each other and how these sites are organized relative to the remainder of the genome.

**(ii) Nuclear Speckle preference.** Interestingly, we found that regions that are closer to the nuclear speckles are strongly associated with high levels of transcriptional activity (Figure 6C). In particular, we found a strong correlation between RNA polymerase II occupancy and nascent RNA transcription levels (measured with GRO-seq^91^) within a genomic region and its distance to a nuclear speckle (Spearman ρ = 0.88 and ρ = 0.75, respectively, Figure 6C, S6D). In fact, the more highly transcribed a genomic region is, the closer it is to a nuclear speckle. However, transcription alone does not explain distance to the nuclear speckles because genomic DNA regions that are not transcribed but are contained within highly transcribed gene dense regions also tend to be closer to the nuclear speckle (Figure 6D). Conversely, highly transcribed genes that are present within otherwise inactive regions tend to be farther from the nuclear speckle (Figure 6D). These results suggest that the density of RNA polymerase II transcription in a genomic neighborhood acts to drive organization around nuclear speckles. This observation may explain the previous observations that gene-dense actively transcribed regions can “loop out” away from the core chromosome territory and can localize proximal to nuclear speckles^16,53,92,93^.

Together, our results demonstrate that preferential association with these nuclear bodies acts to organize the overall packaging of genomic DNA in the nucleus such that regions on different chromosomes with similar functional activity (i.e. transcription levels) can often be closer to each other than to regions on the same chromosome that differ in functional activity (Figure 7).

**Figure 7.**
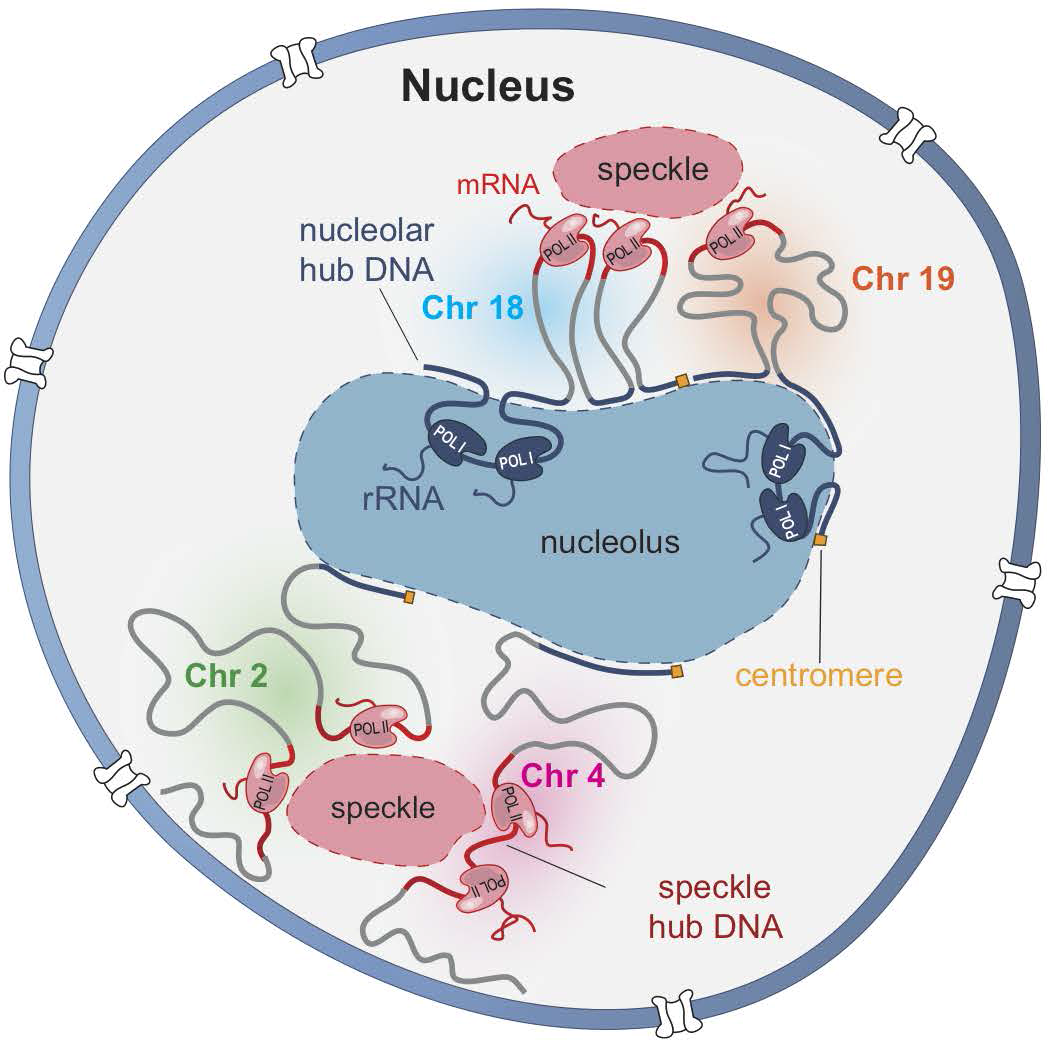
Model for how higher-order inter-chromosomal hubs shape 3-dimensional genome organization in the nucleus. DNA organization in the nucleus is shaped around nuclear bodies. Specifically, transcriptionally inactive regions of chromosomes containing ribosomal DNA genes (e.g., chr18 and chr19, blue) as well as centromere-proximal inactive regions on all chromosomes (e.g., chr2 and chr4, yellow box) can organize around the nucleolus. Transcriptionally active regions with high concentrations of RNA polymerase II co-localize on the periphery of nuclear speckles (red) and preferentially localize in proximity of each other. In this way, regions on different chromosomes that have shared transcriptional states can localize closer to each other around nuclear bodies compared to other regions on the same chromosome.

## DISCUSSION

Our results reconcile two distinct perspectives of nuclear structure into a model that explains how the 3-dimensional structure of the genome is packaged in the nucleus. Specifically, our results uncovered two highly interconnected networks of higher-order inter-chromosomal interactions that are arranged around nuclear bodies and act as organizing centers of overall genome packaging in the nucleus. In addition to the specific nuclear bodies identified here, other nuclear structures including more specialized and smaller structures, may also be involved in shaping DNA structure. Interestingly, the tight relationship between transcriptional status in a region and organization near the nuclear speckles suggests that organization around the nuclear speckle is a highly dynamic and transcriptionally-dependent process. These results contrast with previous views of genome organization that were centered around chromosome territories, a structural feature that is invariant to changes in gene expression^16,24,28^. Together, these results lead to a new picture of genome organization where regions across chromosomes organize around specific nuclear bodies to shape the overall 3-dimensional organization of the nucleus in a highly regulated and dynamic manner (Figure 7).

While the precise functional role of this spatial organization remains to be determined, spatial segregation into different regions of the nucleus may enable more efficient regulation by segregating regulatory factors into regions of high local concentration within the nucleus. For example, this organization may act to enrich repressive complexes near repressed genes, while ensuring that they are depleted near active genomic regions, and vice versa. Specifically, while organization of DNA near nuclear speckles does not appear to impact transcription of individual genes^48,51^, it may provide other regulatory advantages such as by increasing the efficiency of post-transcriptional mRNA processing by concentrating splicing and processing factors, which are enriched in the nuclear speckles, near actively transcribed genes.

More generally, SPRITE represents a powerful new framework for spatial mapping because it provides genome-wide data that is highly analogous to microscopy and can be used to explore large numbers of high resolution combinatorial interactions that occur simultaneously in 3-dimensional space within individual cells. Beyond its current applications, SPRITE can be extended to include direct measurements of RNA and incorporate protein localization^94,95^ relative to the higher-order 3-dimensional genome structure. These applications will enable exploration of previously inaccessible questions regarding the relationship between 3-dimensional genome structure and gene regulation within the nucleus. For example, these approaches will be valuable in directly mapping genome structure relative to other nuclear bodies, structures, and compartments at high resolution as well as exploring the combinatorial and spatial arrangements of multiple enhancer-promoter interactions and their corresponding transcription levels. Furthermore, by enabling combinatorial and spatial maps of DNA, RNA, and protein in the nucleus, SPRITE will enable new insights into the dynamics of nuclear structure and gene regulation across time.

## Acknowledgements

We thank Frank Alber, Joanna Jachowicz, Aaron Lin, Kathrin Plath, John Rinn, Matt Thomson, Ward G. Walkup, and Barbara Wold for comments on the manuscript and helpful suggestions; current and past members of the Guttman lab for their advice and assistance throughout the project; Sigrid Knemeyer for illustrations; and Brittany Belin for naming the technique. Imaging was performed in the Biological Imaging Facility, with the support of the Caltech Beckman Institute and the Arnold and Mabel Beckman Foundation and technical advice from Andres Collazo and Steven Wilbert. Sequencing was performed at the Caltech Millard and Muriel Jacobs Genetics and Genomics Laboratory with assistance from Igor Antoshechkin. Sofia Quinodoz is funded by the Howard Hughes Medical Institute Gilliam Fellowship and NSF GRFP Fellowship. Prashant Bhat is funded by NIGMS T32 GM008042 and the UCLA-Caltech Medical Scientist Training Program. This work was funded by grants from the NIH 4D Nucleome Project (U01 DA040612 and U01 HL130007), NHGRI Genomics of Gene Regulation program (U01 HG007910 to M. Garber), the New York Stem Cell Foundation, an NIH Director’s Early Independence Award (DP5OD012190), the Edward Mallinckrodt Foundation, Sontag Foundation, Searle Scholars Program, Pew-Steward Scholars program, and funds from the California Institute of Technology. Mitchell Guttman is a New York Stem Cell Foundation – Robertson Investigator.

## SUPPLEMENTAL FIGURE LEGENDS

**Figure S1.**
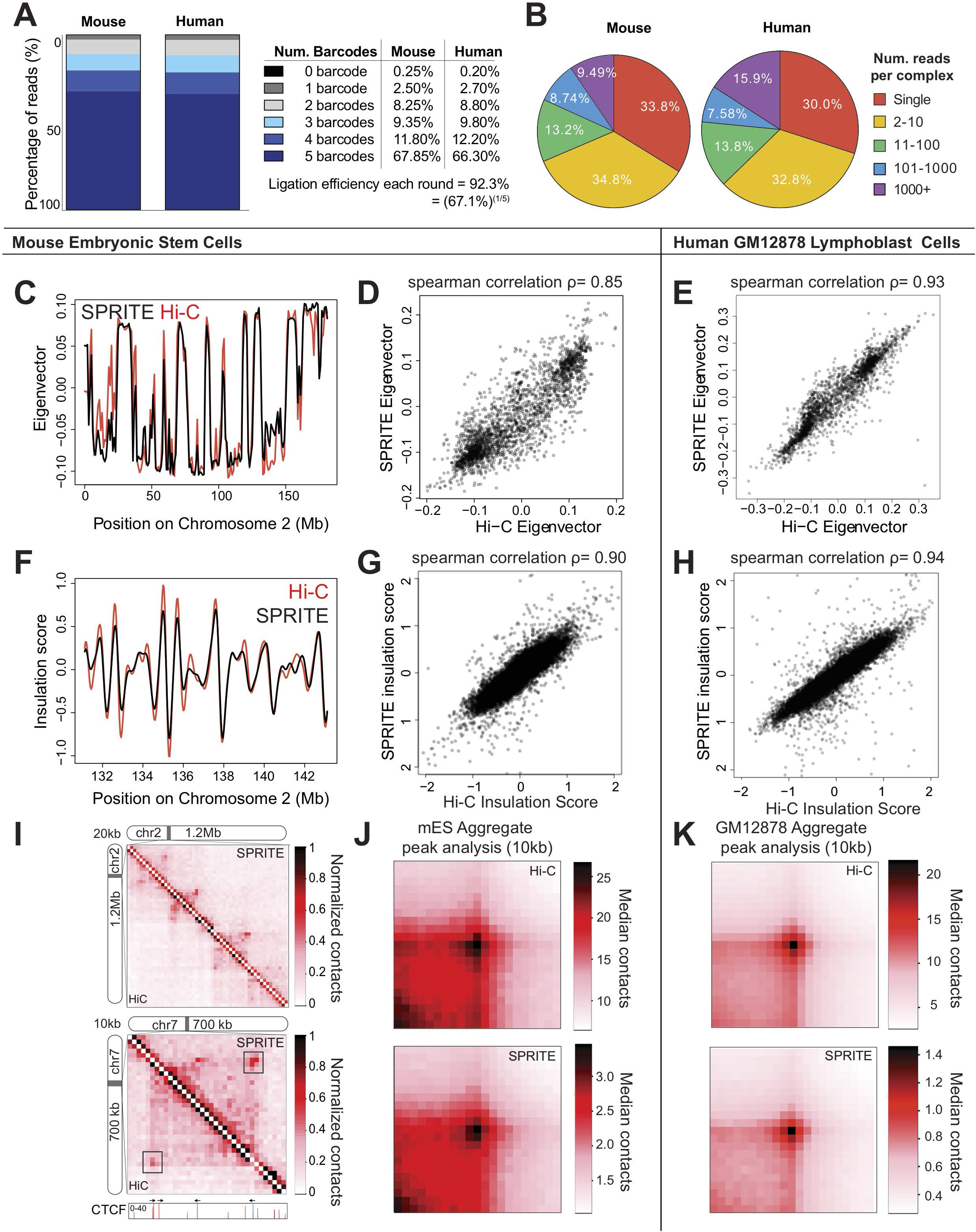
SPRITE accurately recapitulates genome structure measured by Hi-C. (A) Distribution of reads containing 0, 1, 2, 3, 4, 5 barcodes for mouse (left) and human (right) experiments. The estimate for ligation efficiency each round is determined by taking the 5^th^ root of the fraction of reads with all 5 barcodes. (B) Distribution of SPRITE cluster sizes for mouse (left) and human (right) experiments. The number of reads was calculated for different SPRITE cluster sizes (1, 2-10, 11-100, 101-100 and over 1001) and reported as the percentage of total reads. (C) Compartment eigenvector for mouse chromosome 2 calculated using SPRITE (black) and Hi-C (red) contact maps from mouse ES cells binned at 1Mb resolution. Positive and negative values correspond to the A and B compartments, respectively. (D) Genome-wide correlation between compartment eigenvectors calculated using SPRITE (y axis) and Hi-C (x axis) contact maps from mouse ES cells binned at 1Mb resolution. (E) Genome-wide correlation between compartment eigenvectors calculated using SPRITE (y axis) and Hi-C (x axis) contact maps from human GM12878 cells binned at 1Mb resolution. (F) Insulation score profile for a region on mouse chromosome 2 (shown in Figure 1D) calculated using SPRITE (black) and Hi-C (red) contact maps from mouse ES cells binned at 40kb resolution. Local minima correspond to boundary regions. (G) Genome-wide correlation between insulation scores calculated using SPRITE (y axis) and Hi-C (x axis) contact maps from mouse ES cells binned at 40kb. (H) Genome-wide correlation between insulation scores calculated using SPRITE (y axis) and Hi-C (x axis) contact maps from human GM12878 cells binned at 40kb. (I) Examples of SPRITE and Hi-C contact maps binned at 20kb resolution (top) and 10kb resolution (bottom) showing chromatin loop interactions. CTCF ChIP-seq peaks are shown according to their positive (red) or negative (blue) motif orientation. (J) Aggregate peak analysis heatmaps for Hi-C (top) and SPRITE (bottom) in mouse ES cells binned at 10kb resolution. 1493 loops obtained from ^23^ were used in this analysis. Heatmaps show the median contact map values for each pair of 10kb bins in regions +/− 200kb of the loops. (K) Aggregate peak analysis heatmaps for Hi-C (top) and SPRITE (bottom) in human GM12878 cells binned at 10kb resolution. 5789 loops obtained from Rao *et al.*^23^ were used in this analysis. Heatmaps show the median contact map values for each pair of 10kb bins in regions +/- 200kb of the loops.

**Figure S2.**
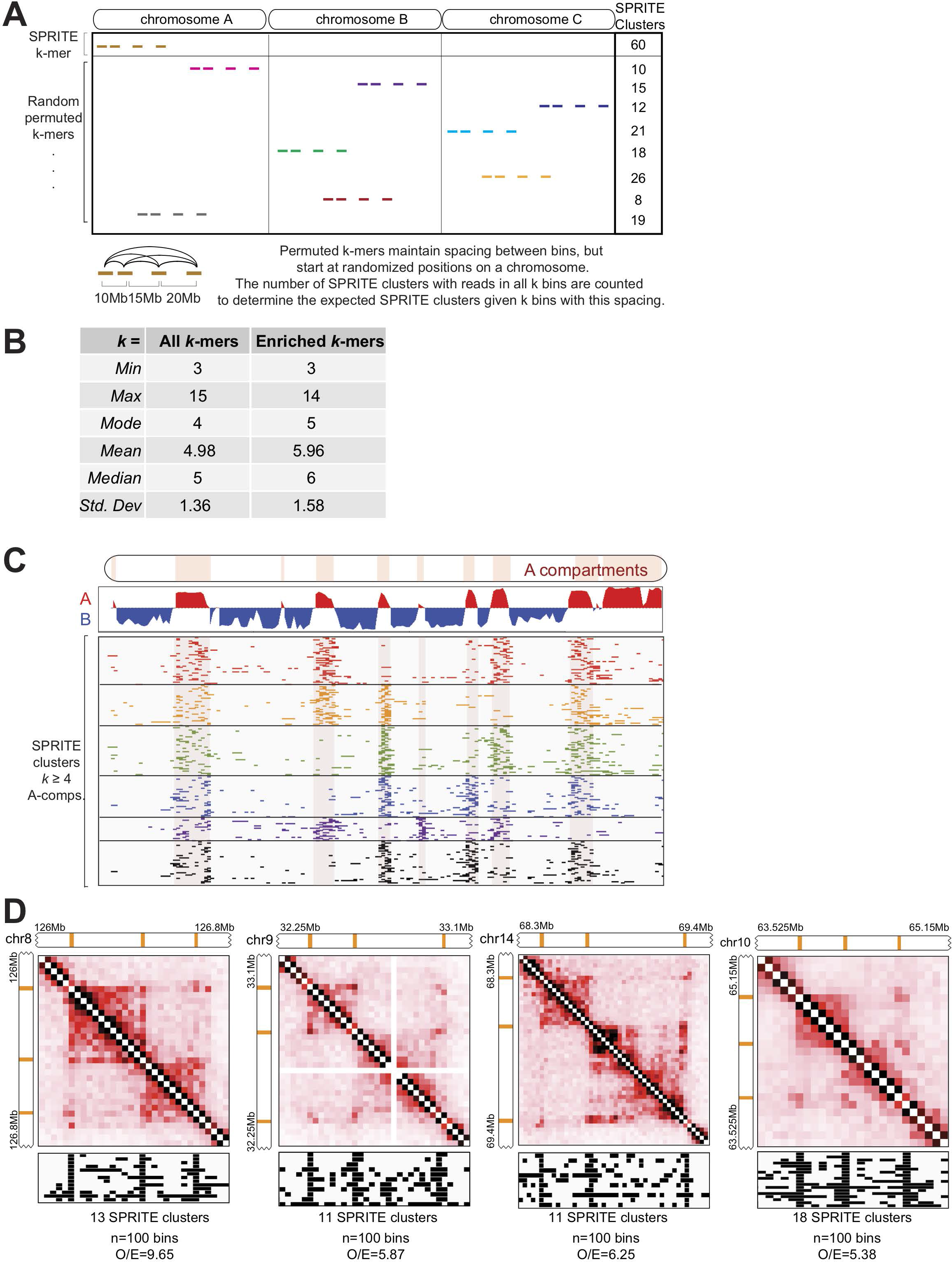
SPRITE measures higher-order interactions across known genome structures. (A) Cartoon representation of the method for identifying enriched higher-order *k*-mer frequencies. Given an observed *k*-mer, randomly permuted *k*-mers are sampled by randomizing the location of the *k*-mer while preserving the spacing between its reads. The observed frequency is then normalized by the expected frequency derived from the randomly permuted *k-*mers. (B) Statistics of the number of *k* bins observed in all enumerated (left) and enriched (right) *k*-mers in mES cells at 1Mb resolution. Enriched *k*-kmers are defined as those that are observed in at least 5 independent SPRITE clusters, occur >4-fold more frequently than the average of the permuted *k*-mers, and are observed at a frequency that is more than 90% of the permuted structures. (C) Example SPRITE clusters spanning four A compartment regions on mouse chromosome 2. The compartment eigenvector showing A (red) and B (blue) compartments is shown on top. Rows correspond to individual SPRITE clusters and colored lines denote 1Mb bins with at least 1 read. Each colored group represents a different combination of four A compartment regions interacting across several SPRITE clusters (max cluster size n=20 1Mb bins). Red boxes demarcate the A compartment regions. Enrichments compared to randomly permuted *k*-mers that spanning 4 or more A compartment regions in mES cells are listed in **Table S2**. (D) Four examples of 3-way interactions between 3 loop anchors in human GM12878 cells. SPRITE contact maps are shown at 25kb resolution above the individual SPRITE clusters that contain all three loop anchors.

**Figure S3.**
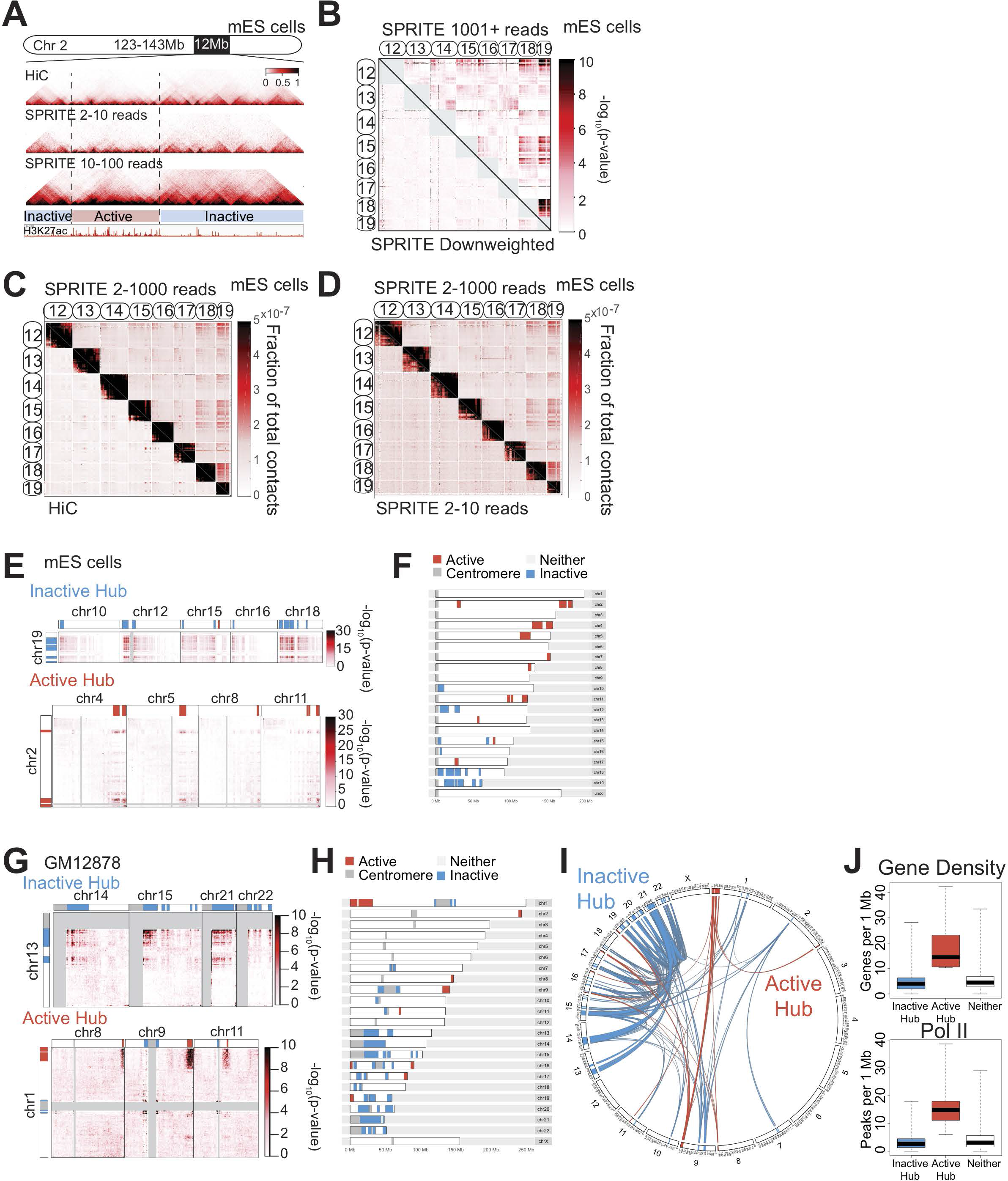
SPRITE identifies long-range intra-chromosomal interactions and hubs of inter-chromosomal interactions. (A) Comparison of Hi-C contact map (top) with SPRITE contact maps based on SPRITE clusters with 2 to 10 reads (middle) and 10 to 100 reads (bottom) in a 12Mb region on mouse chromosome 2 binned at 40kb resolution. H3K27ac ChIP-seq signal from mouse ES cells is shown below. (B) Inter-chromosomal contacts between chromosomes 12 through 19 in mES cells binned at 1Mb resolution. Interaction p-values are shown for SPRITE clusters with 1001+ reads (upper diagonal) and for SPRITE contacts that have been down-weighted by cluster size (lower diagonal). *p*-values were calculated based on inter-chromosomal interaction frequencies and are shown in units of −log_10_ (*p*-value). (C) Comparison of SPRITE (upper diagonal) and Hi-C (lower diagonal) inter-chromosomal contacts between chromosomes 12 through 19 in mES cells binned at 1Mb resolution. SPRITE contacts are based on clusters with 2 to 1000 reads. Values shown are the fraction of total contacts. (D) Comparison of inter-chromosomal contacts between chromosomes 12 through 19 based on SPRITE clusters with 2 to 1000 reads (upper diagonal) or 2 to 10 reads (lower diagonal) in mES cells binned at 1Mb resolution. Values shown are the fraction of total contacts. (E, G) Examples of inter-chromosomal interactions that comprise the Inactive Hub (top, blue) and Active Hub (bottom, red) in mES and GM12878 cells, respectively. Inactive and active hub regions are colored as blue and red, respectively. Interaction p-values were calculated using unweighted contact frequencies from SPRITE clusters with 2 to 1000 reads and are shown in units of –log_10_ (*p-*value). (F, H) Ideogram showing inactive hub (blue) and active hub (regions) on each mouse and human chromosome, respectively. Centromere regions are demarcated in grey. (I) Circos diagram of significant inter-chromosomal interactions in GM12878 cells. Interactions are identified from unweighted inter-chromosomal contact maps in SPRITE clusters containing 2 to 1000 reads. Interactions with *p*-values less than 10^−8^ are shown. Two distinct networks of noninteracting connections are shown in blue (inactive hub) and red (active hub). (J) Box plots showing the distribution of gene density (top) and RNA polymerase II (Pol II) occupancy (bottom) for regions in the inactive hub, nucleolar hub, or neither hub in GM12878 cells. Gene density is calculated as genes per 1Mb and Pol II occupancy is calculated as the number of ChIP-seq peaks per 1Mb.

**Figure S4.**
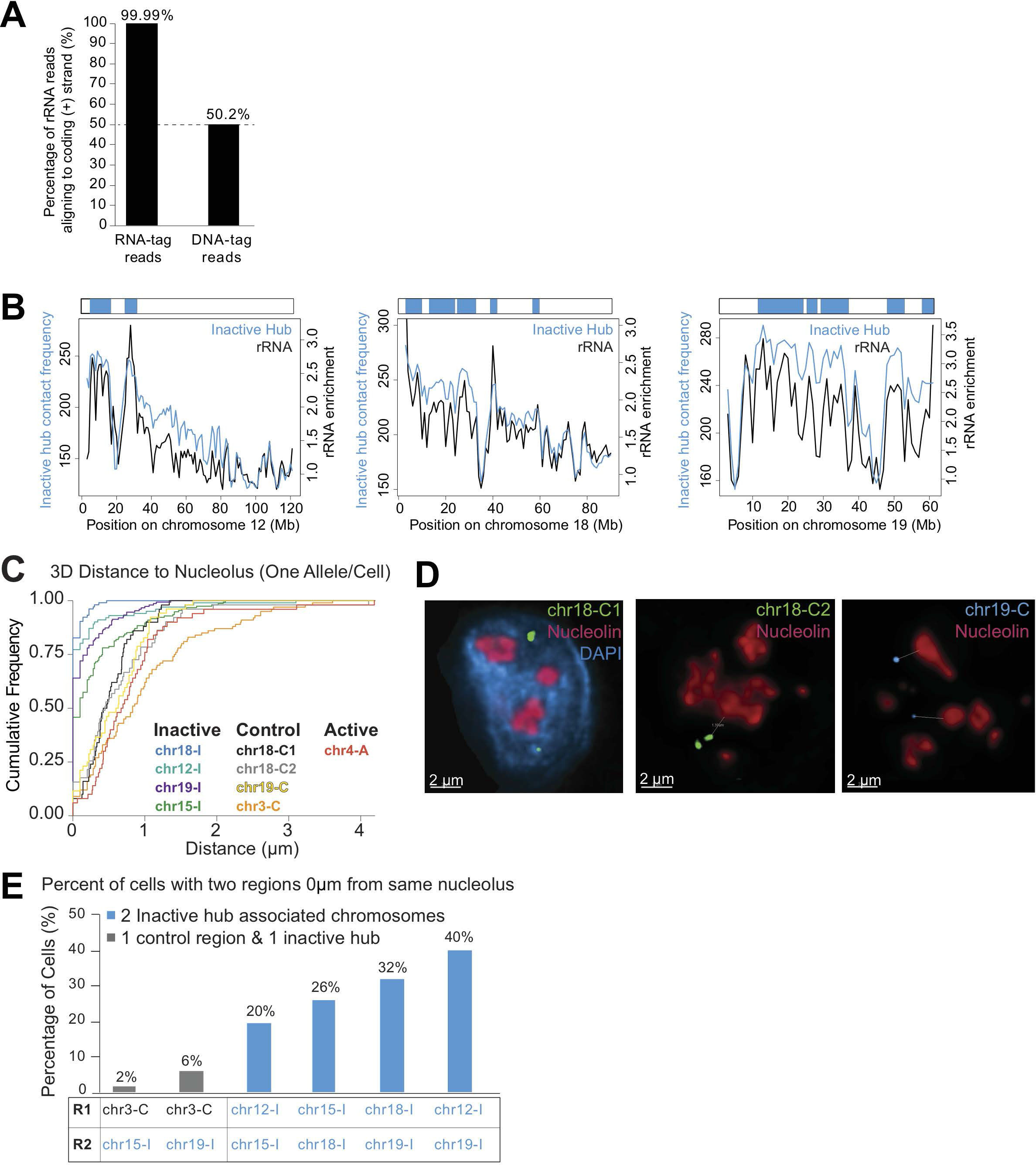
Genomic DNA in the inactive hub is organized around the nucleolus. (A) Comparison of coding strand bias for reads tagged with an RNA- or DNA- specific adaptor that aligned to ribosomal RNA in the RNA-DNA SPRITE maps. (B) Comparison of ribosomal RNA localization (black) and average inter-chromosomal contact frequency with regions in the inactive hub (blue) binned at 1Mb resolution for mouse chromosomes 12, 18 and 19. Inactive hub regions are shown in blue at the top of each plot. (C) Cumulative distributions for the 3D distances 8 probe regions and the most proximal nucleolus (only the distance of the nearest allele per cell is shown). Four probe regions in the inactive hub are shown in blue, green, teal, and purple. Four control probe regions not in the inactive hub are shown in black, grey, orange, and yellow. One active hub probe region is shown in red. (D) Example images from immunofluorescence for nucleolin (red) combined with DNA FISH for three different DNA FISH control region probes (chr18-C1, chr18-C2 and chr19-C). (E) Comparison of the frequency of co-localization of at least one allele for six different pairs of DNA FISH probes at the same nucleolus. Pairs of probes with one control region and one inactive hub region are shown in grey and pairs of two inactive hub region probes are shown in blue.

**Figure S5.**
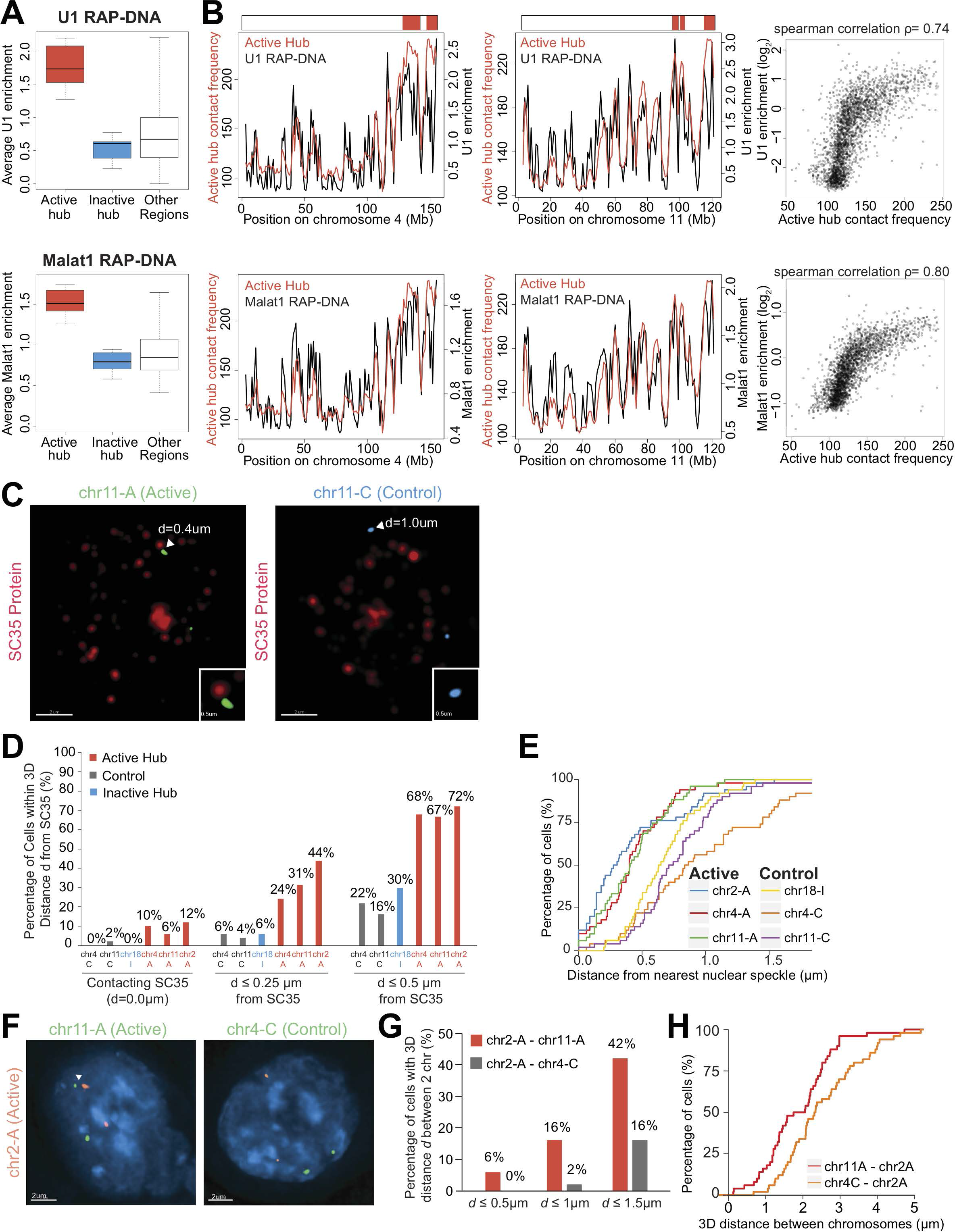
Genomic DNA regions in the active hub are organized around the nuclear speckles. (A) Box plots showing the distribution of U1 RAP-DNA enrichment (top) and Malat1 RAP-DNA enrichment (bottom) from Engreitz *et al.*^79^ for regions in the active hub, inactive hub or neither. (B) Comparison of U1 or Malat1 RAP-DNA (black) enrichment with average inter-chromosomal contact frequency with regions in the active hub binned at 1Mb resolution for mouse chromosomes 4 (left) and 11 (middle). Genome-wide correlation plots are shown on the right. (C) Example images from immunofluorescence for SC35 (red) combined with DNA FISH for two different probe regions. Inset images show the region around the arrowhead. These cells were not stained for DAPI due to the emission of the DNA FISH probes used, and therefore lack a nuclear signal. (D) Comparison of proximity to the most proximal nuclear speckle for 6 different probe regions, including 3 control regions (grey/blue) and 3 active hub regions (red). Proximity to nuclear speckles is calculated for each probe region as the percent of cells with at least 1 allele within 0 μm (left), 0.25 μm (middle) or 0.5 μm (right) of an SC35 speckle. (E) Cumulative distributions for the 3D distances of 6 probe regions and the most proximal speckle (only the distance of the nearest allele per cell is shown). Three probe regions in the active hub are shown in blue, red, and green. Three control probe regions not in the active hub are shown in yellow, orange, and purple. (F) Example images from DNA FISH for two pairs of probe regions. The left image shows two active hub region probes (chr2-A and chr11-A) and the right image shows one active hub region probe (chr2-A) and a control region probe (chr4-C). (G) Comparison of the percent of cells where two probes are within a given distance (0.5 μm, 1.0 μm and 1.5 μm) for two active hub probes (red) and an active/control pair (grey). (H) Cumulative distributions for the 3D distances between two pairs of probe regions: chr2-A and chr11-A (red), and chr2-A and chr4-C (orange) (only the distance of the pair of the closest alleles are down).

**Figure S6.**
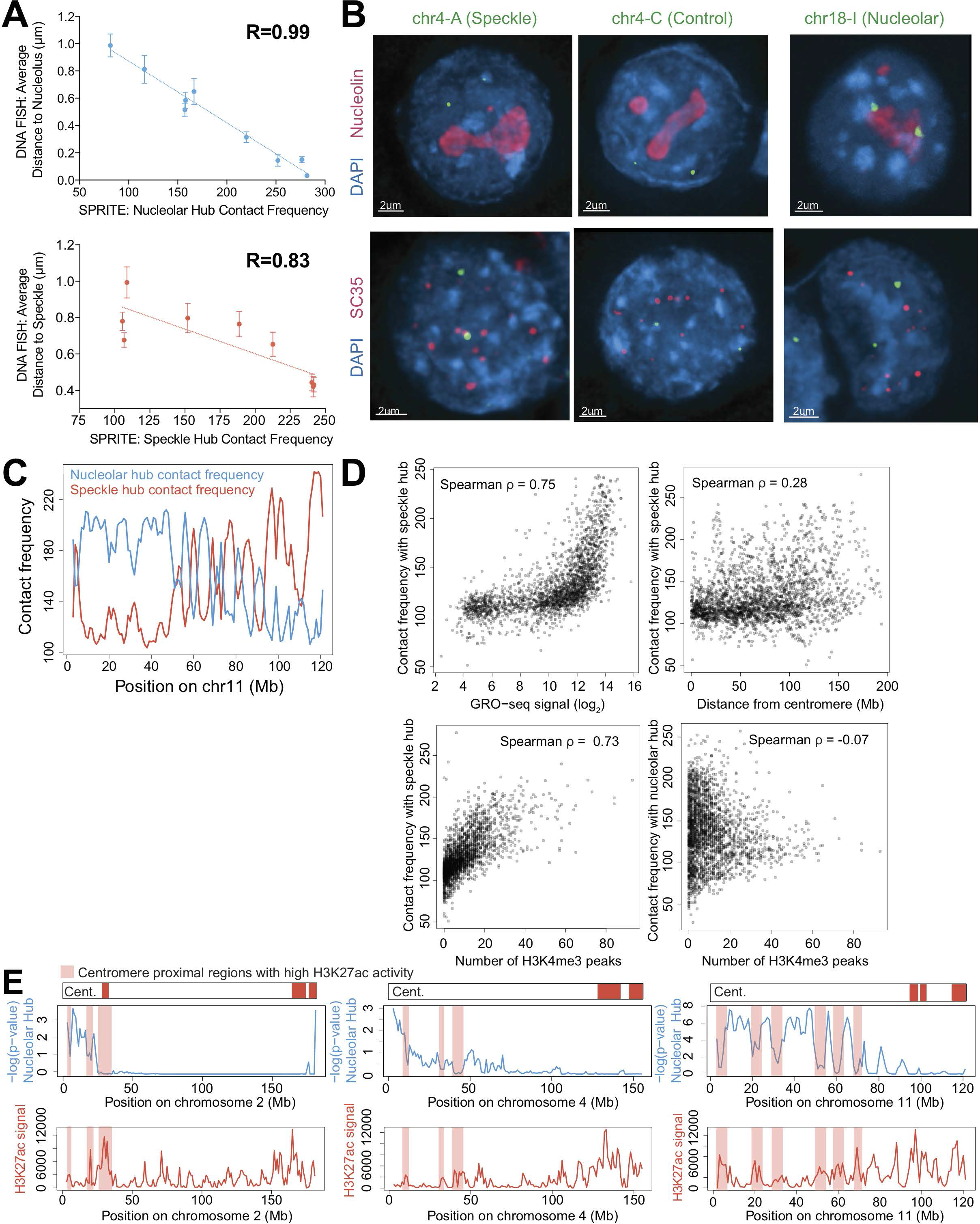
Preferential organization relative to the nucleolus and nuclear speckles shapes the overall organization of genomic DNA in the nucleus. (A) Comparison of SPRITE contact frequencies with the nucleolar hub (left) or speckle hub (right) regions with DNA FISH average 3D distances from nucleoli (left) or nuclear speckles (right). Average 3D distances are measured from DNA FISH and IF experiments using SC35 and nucleolin as markers for nuclear speckles and nucleoli, respectively. The closest allele in each cell is used to measure the 3D distance to each nuclear body. (B) Examples images from immunofluorescence combined with DNA FISH showing three regions with distinct association patterns with nucleoli (top row) and nuclear speckles (bottom row). chr4-A (left) associates with nuclear speckles but not nucleoli, chr4-C (middle) associates with neither, and chr18-I (right) associates with nucleoli but not nuclear speckles. (C) Average inter-chromosomal contact frequency to regions in the nucleolar hub (blue) or speckle hub (red) for all 1Mb regions on mouse chromosome 11. These inter-chromosomal contact frequencies are computed from unweighted interactions using SPRITE clusters containing 2 to 1000 reads. (D) Comparison of average inter-chromosomal contact frequencies with the speckle hub and nascent transcription levels (GRO-seq signal from the Lis lab^91^), linear distance from centromeres, and number of H3K4me3 peaks (ENCODE). Comparison of average SPRITE inter-chromosomal contact frequencies with the nucleolar hub and H3K4me3 peaks. (E) Examples of decreased nucleolar hub interactions in centromere-proximal regions with high levels of transcriptional activity (light red box). Although there is a general trend of centromere-proximal regions to interact with the nucleolar hub, highly active regions near centromeres are depleted in nucleolar hub interactions (blue). H3K27ac ChIP-seq (ENCODE) signal (red) is shown below.

**Figure S7.**
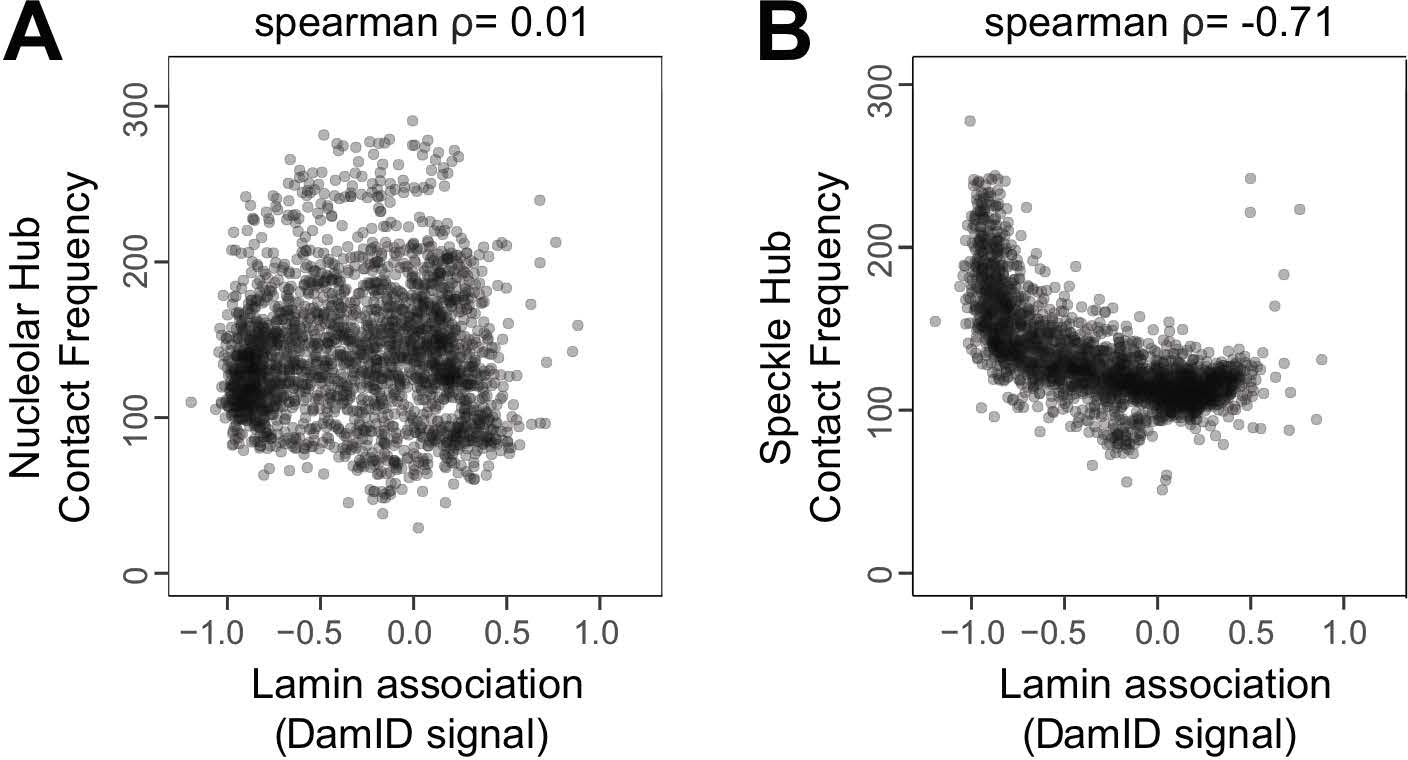
Nucleolar association is not correlated with lamin association, but speckle hub association is inversely correlated with lamin association. (A) Comparison between DamID signal from Meuleman *et al.*^96^ and nucleolar hub contact frequencies in mES cells. (B) Comparison between DamID signal and speckle hub contact frequencies in mES cells.

## SUPPLEMENTAL TABLES

**Table S1. Enriched** ***k-mers*** **in mES cells.** All *k-mers* at 1 megabase resolution that were observed in at least 5 independent SPRITE clusters, at an observed frequency that exceeded 90% of the random permutations, and occurred at least than 4-times more frequently than the average of the permutated regions. Each *k-*mer was randomly permuted in a manner that preserves genomic distance between all regions within the *k*-mer.

**Table S2.** ***k*****-mers spanning three or four A compartment regions on individual chromosomes in mES cells.** All *k*-mers observed in SPRITE clusters that contain reads from three or four different A compartment regions that span at least 100Mb within an individual chromosome. These *k-*mers were randomly permuted 100 times in a manner preserving genomic distance between these regions.

**Table S3.** ***k*****-mers spanning three or more nucleolar hub regions on different chromosomes.** Enrichments of *k-*mers containing at least three distinct nucleolar hub regions that were present on different chromosomes compared to inter-chromosomal *k*-mers that preserve the size of the hub regions and were randomly permuted 100 times across 3 different chromosomes.

**Table S4.** ***k*****-mers spanning three or more active hub regions on different chromosomes.** Enrichments of of *k-*mers containing at least three distinct active hub regions that were present on different chromosomes compared to inter-chromosomal *k*-mers that preserve the size of the hub regions and were randomly permuted 100 across 3 different chromosomes.

**Table S5. Sequences of SPRITE barcoded adaptors used.** Barcoded DPM, Odd, Even, Terminal adaptors that were used in this study and their sequences are listed in this table.

## SUPPLEMENTAL MOVIE

**Movie S1. Nucleolar hub regions co-associate around the same nucleolus in an individual cell.** Nucleolar hub regions on chromosome 15 (orange) and 18 (green) co-associate around the same nucleolus (red), marked by Nucleolin using immunoflourescence. DAPI is used to stain DNA in the nucleus.

## MATERIALS AND METHODS

### Cell culture and lines used in analysis

Mouse ES cell lines were cultured in serum-free 2i/LIF medium and maintained at an exponential growth phase as previously described^97–99^. SPRITE DNA-DNA maps were generated in female ES cells (F1 2-1 line, provided by K. Plath), an F1 hybrid wild-type mouse ES cell line derived from a 129 × *castaneous* cross. SPRITE RNA-DNA maps were generated in the pSM33 ES cell line (provided by K. Plath), a male ES cell line derived from the V6.5 ES cell line, which expresses Xist from the endogenous locus under the transcriptional control of a tet-inducible promoter and the Tet transactivator (M2rtTA) from the Rosa26 locus. We induced Xist expression in these cells using doxycycline (Sigma, D9891) at a final concentration of 2ug/ml for 6-24hrs.

Human GM12878 cells, a lymphoblastoid cell line obtained from Coriell Cell Repositories, were cultured in RPMI 1640 (Gibco, Life Technologies), 2mM L-glutamine, 15% fetal bovine serum, and 1x penicillin-streptomycin and maintained at 37**°**C under 5% CO_2_. Cells were seeded every 3-4 days at 200,000 cells/ml in T25 flasks, maintained at an exponential growth phase, and passaged or harvested before reaching 1,000,000 cells/ml.

### Split-Pool Recognition of Interactions by Tag Extension (SPRITE)

***Crosslinking.*** Cells were crosslinked in a single-cell suspension to ensure that we obtain individual crosslinked nuclei rather than crosslinked colonies of cells. For GM12878 lymphoblast cells, which are grown in suspension, cells were spun and media was removed prior to crosslinking. For mouse ES cells, which are adherent, cells were trypsinized to remove from plates prior to crosslinking in suspension. Specifically, 5mL TVP (1mM EDTA, 0.025% Trypsin, 1% Sigma Chicken Serum; pre-warmed at 37C) was added to each 15cm plate, then rocked gently for 3-4 minutes until cells start to detach from the plate. Afterwards, 25mL wash solution (DMEM/F-12 supplemented with 0.03% Gibco BSA Fraction V, pre-warmed at 37C) was added to each plate to inactivate the trypsin. Cells were lifted into a 15 or 50ml conical tube, pelleted at 330g for 3 min, and then washed in 4mL of 1X PBS per 10mL cells. During all crosslinking steps and washes, volumes were maintained at 4mL of buffer/crosslinking solution per 10M cells. After pelleting, cells were pipetted to disrupt clumps of cells and crosslinked in suspension with 4mL of 0.5M disuccinimidyl glutarate (DSG, Pierce) dissolved in 1X PBS for 45 minutes at room temperature. DSG was removed, cells were pelleted (as above) and washed with 1X PBS, and 3% formaldehyde (FA Ampules, Pierce) in 1X PBS was added to cells for 10 minutes at room temperature. Formaldehyde was immediately quenched with addition of 200μl of 2.5M Glycine per 1mL of 3%FA solution, cells were pelleted, formaldehyde was removed, and cells were washed in ice cold 1X PBS + 0.5% BSA, 3 times. Cells were then aliquoted into tubes containing 10 million cells each, pelleted, supernatant removed, and flash frozen in liquid nitrogen and stored in −80°C until lysis.

***Chromatin Isolation.*** Crosslinked cell pellets (10 million cells) were lysed using the nuclear isolation procedure as previously described by Blecher-Gonen *et al.*^100^ with minor modifications. Specifically, cells were incubated in 1mL of nuclear isolation buffer 1 (50mM Hepes pH 7.4, 1mM EDTA pH 8.0, 1mM EGTA pH 8.0, 140mM NaCl, 0.25% Triton-X, 0.5% NP-40, 10% Glycerol, 1xPIC) for 10 minutes on ice. Then cells were pelleted at 850g for 10 minutes at 4°C. Supernatant was removed and 1ml of Lysis Buffer 2 (50mM Hepes pH 7.4, 1.5mM EDTA, 1.5mM EGTA, 200mM NaCl, 1xPIC) was added and incubated for 10 minutes on ice. Nuclei were obtained after pelleting and supernatant removed (as above) and 550ul of Lysis Buffer 3 (50mM Hepes pH 7.4, 1.5mM EDTA, 1.5mM EGTA, 100mM NaCl, 0.1% Sodium deoxycholate, 0.5% NLS, 1xPIC) was added and incubated for 10 minutes on ice prior to sonication.

***Chromatin Digestion.*** After nuclear isolation, we digested chromatin by sonicating the nuclear pellet using a Branson needle-tip sonicator at 4°C for a total of 1 minute at 4-5 watts (pulses of 0.7 seconds on, followed by 3.3 seconds off). DNA was further digested using 2 - 6 uL of TurboDNAse (Ambion) per 10ul of sonicated lysate (equivalent to ∼200,000 cells), in 10x DNase Buffer (200mM Hepes pH 7.4, 1M NaCl, 0.5% NP-40, 5mM CaCl_2_, 25mM MnCl_2_) at 37°C for 20 min. Concentrations of DNase were optimized to obtain DNA fragments of approximately 150bp-1000bp, which is needed for sequencing these fragments. DNAse activity was quenched by adding 10mM EDTA and 5mM EGTA.

***Estimating molarity.*** After DNAse digestion, approximately 20μl of DNAsed lysate was reverse crosslinked in 80ul of 1x Proteinase K buffer (20mM Tris pH 7.5, 100mM NaCl, 10mM EDTA, 10mM EGTA, 0.5% Triton-X, 0.2% SDS) with 8ul Proteinase K (NEB) at 65°C overnight. The DNA was purified using Zymo DNA Clean and Concentrate columns per the manufacturer’s specifications with minor adaptations, such as binding to the column with 7x Binding buffer to improve yield. The molarity of DNA was calculated by measuring the concentration of DNA with the Qubit Fluorometer (HS dsDNA kit) and the sizes were estimated using the Agilent Bioanalyzer (HS DNA kit).

***NHS bead coupling.*** We used these numbers to calculate the total number of DNA molecules per microliter of lysate. We coupled the lysate to NHS-activated magnetic beads (Pierce) overnight at 4°C in 1ml of 1X PBS + 0.1% SDS rotating on a HulaMixer Sample Mixer (Thermo). Specifically, we coupled 1x10^10^ to 1.75mls of beads (mouse) and 5x10^10^ molecules to 2 mls of beads (human). We obtain roughly 50% coupling efficiency of molecules to the beads, which effectively halves the ratio of molecules coupled per bead. This coupling ratio was selected to ensure that most beads contained less than 0.125 to 0.25 complex per bead to reduce the probability of simultaneously coupling multiple independent complexes to the same bead, which would lead to their association during the split-pool barcoding process. At this loading concentration, we find that <0.25% of SPRITE clusters are inter-species and <0.5% of contacts contain any spurious pairing of human and mouse fragments that arise due to bead coupling (see Figure M1).

After coupling lysate to NHS beads overnight, we quench the beads in 1mL of 0.5M Tris pH 8.0 for 1 hour at 4C rotating on a HulaMixer. We then wash the beads 4 times at 4C in 1mL of Modified RLT Buffer (1x Buffer RLT supplied by Qiagen with added 10mM Tris pH 7.5, 1mM EDTA, 1mM EGTA, 0.2% NLS, 0.1% Triton-X, 0.1% NP-40) for 3 minutes each. Next, beads are washed in 1mL of SPRITE Wash Buffer (1x PBS, 5mM EDTA, 5mM EGTA, 5mM DTT, 0.2% Triton-X, 0.2% NP-40, 0.2% Sodium deoxycholate) twice at 50C and once at room temperature for 5 minutes each. These washes remove any material that is not covalently attached to the beads. Prior to performing all enzymatic steps, buffer is exchanged on the beads through 2 rinses using 1mL of 1x Detergent Buffer (20mM Tris pH 7.5, 50mM NaCl, 0.2% Triton-X, 0.2% NP-40,0.2% Sodium deoxycholate). These detergents are used throughout the protocol to prevent bead aggregation, which could result in spurious interactions. Because the crosslinked complexes are immobilized on NHS magnetic beads, we can perform several enzymatic steps by adding buffers and enzymes directly to the beads and performing rapid buffer exchange between each step on a magnet. All enzymatic steps were performed with shaking at 1200 rpm (Eppendorf Thermomixer) to avoid bead settling and aggregation, and all enzymatic steps were inactivated with addition of 0.5-1mL Modified RLT Buffer to NHS beads.

***DNA Repair.*** We then repair the DNA ends to enable ligation of tags to each molecule. Specifically, we blunt end and phosphorylate the 5’ ends of double-stranded DNA using two enzymes. (i) T4 Polynucleotide Kinase (NEB) treatment is performed at 37°C for 1 hour, the enzyme is quenched using 1mL Modified RLT buffer, and then buffer is exchanged twice using washes of 1mL 1x Detergent Buffer to beads at room temperature. Next, the NEBNext End Repair Enzyme cocktail, containing T4 DNA Polymerase and T4 PNK, and mastermix is added to beads and incubated at 20°C for 1 hour, inactivated and buffer exchanged as specified above. DNA was then dA-tailed using the Klenow fragment (5’-3’ exo-, NEBNext dA-tailing Module) at 37°C for 1 hour, and inactivated and buffer exchanged as specified above.

***Split-pool ligation.*** The beads were then repeatedly split-and-pool ligated over 5 rounds with a set of DNA phosphate Modified (DPM), “Odd”, “Even” and “Terminal” tags (see **SPRITE Tag Design** below for details). The DPM tag is ligated by an “Odd” tag. The “Odd” and “Even” tags were designed so that they can be ligated to each other over multiple rounds, such that after Odd is ligated, then Even ligates the Odd tags, and then Odd can ligate the Even tags. This can be repeated such that the same 2 plates of tags can be used over multiple rounds of split-pool tagging without self-ligation of the adaptors to each other. Finally, a set of barcoded terminal tags are ligated at the end to attach an Illumina sequence for final library amplification. In this study, we performed 5 rounds total of split-and-pool ligation in the following order: DPM, Odd, Even, Odd, and Terminal tag. Over each round, the samples are split across a 96 well plate in 6.5ul per well of 1x SPRITE Detergent Buffer to prevent aggregation of beads, which would result in spurious interactions. Each plate contained 2.4ul of 96 different tags at a concentration of 45uM. 10ul of 2x Instant Sticky End Ligation Master Mix (NEB) and 1.1ul of Ultra Pure H20 (Invitrogen) was added to each well of the 96 well plate, for a final concentration of 1x Instant Sticky End Ligation Master Mix per well. All ligations were performed at 20°C for 1 hour with shaking at 1600 rpm for 30 seconds every 5 min. Following every round of split-pool ligation, we inactivated the ligase via addition of 60ul of Modified RLT Buffer to every well, which prevents spurious ligation of tags in the pooled tube. The sample was then pooled into a single 1.7mL tube. After removing Modified RLT Buffer from the beads, remaining free tags were removed by washing the beads in 1mL SPRITE Wash Buffer three times at 45°C for 3 minutes each. We then performed buffer exchange into SPRITE Detergent Buffer by adding 1mL of Buffer and exchanging 3 times.

***Estimating sequencing depth.*** SPRITE interactions are defined based on the sequences that share the same tags. Accordingly, it is essential to sequence all of the barcoded molecules in a complex in order to identify interactions in a sample. Therefore, the number of unique molecules that are sequenced dramatically affects the likelihood of identifying interacting molecules. To address this, we optimized the loading density of our sequencing sample based on the number of unique molecules contained in the sample. Our goal is to load approximately equimolar unique molecules as the number of sequencing reads generated. Specifically, based on our simulations of Poisson sampling, we have found that sequencing with ∼1-3x coverage of reads per the number of unique molecules will ensure that most molecules are sampled. This follows Poisson sampling where 1-1/e^c^ of molecules are sampled at a given *c* coverage. For example, 3x, 2x, and 1x coverage samples approximately 95%, 86%, and 63% of interactions, respectively. In this study, most libraries were sampled with approximately 1.5-2x coverage.

To determine the number of unique molecules in our sample, we measure the amount of material present on beads prior to reverse crosslinking all interactions. To do this, we take an aliquot of the sample and reverse crosslink to elute (as above), cleanup DNA, and PCR amplify for 9-12 cycles. We then measure the molarity using the Qubit and Bioanalyzer (as above). The number of unique molecules prior to PCR is back calculated from a standard curve and adjusted to account for loss during the cleanup. This is used to estimate the number of unique molecules in the sample prior to PCR. In addition to optimizing molarity, because this dilution results in approximately 1% aliquots of the total sample being separately eluted and amplified, this effectively serves as another round of split-pool barcoding as each library is tagged with a unique barcoded Illumina primer. This further reduces the probability that molecules in different clusters obtain the same barcodes.

***Sequencing library generation.*** We ensured that the number of unique molecules loaded (prior to amplification) does not exceed the number of molecules that can be sequenced (∼150-300 million). Aliquots were selected to contain approximately 50-150 million unique molecules, Proteinase K (NEB) digested for 1hr at 50°C in Proteinase K Buffer (20mM Tris pH 7.5, 100mM NaCl, 10mM EDTA, 10mM EGTA, 0.5% Triton-X, 0.2% SDS), and reverse crosslinked overnight at 65°C. DNA was isolated using the Zymo DNA Clean and Concentrator IC. Libraries were amplified using NEB Q5 Hot-Start Mastermix with primers that add the full Illumina adaptor sequence. After amplification, the libraries are cleaned up using 0.7X SPRI (AMPure XP) twice to remove excess adaptors.

***Mapping RNA and DNA simultaneously using SPRITE.*** To map RNA and DNA interactions simultaneously, the SPRITE protocol was performed with the following modifications. (i) Upon coupling of lysate to NHS beads, RNA overhangs caused by fragmentation are repaired by a combination of FastAP treatment (Thermo) and T4 Polynucleoide Kinase (no ATP) at 37°C for 15 min and 1 hr, respectively. RNA was subsequently ligated with a RNA phosphate modified (RPM) tag using NEB ssRNA Ligase 1 (High Concentration) at 20°C for 1 hr, which is designed with a 5’ ssRNA overhang and 3’ dsDNA sticky end for sequential ligation of DNA tags to the RNA (see **Tag design**). (ii) RNA was converted to cDNA using Superscript III (Thermo) using a manganese RT protocol^101^ to promote reverse-transcription through formaldehyde crosslinks on RNA. After cDNA synthesis, cDNA was selectively eluted using RNaseH (NEB) and RNAse Cocktail (Ambion). cDNA was ligated with an unique cDNA tag as previously described,^102^ which serves as a RNA-specific identifier of reads as RNA during sequencing.

Detailed SPRITE protocols are available at www.lncRNA.caltech.edu/SPRITE/

### Human-mouse mixing experiment

Human HEK293T cells and mouse pSM33 cells were crosslinked, lysed, and DNAse digested separately (Figure M1A). The two lysates were then combined at equimolar concentration and coupled to NHS beads at a ratio of 620, 125, 60, 25, 6, 1.2, 0.5, 0.25, 0.125 molecules per NHS bead. Reads were aligned to both hg19 and mm9. All reads aligning to both species were removed, and species-specific reads were used to determine the amount of inter-species contacts. For the experiments in the paper, we selected a coupling efficiency of 0.125 to 0.25 molecules/bead because it provided a small number of spurious contacts while minimizing the number of beads used in the experiment (Figure M1B).

**Figure M1.**
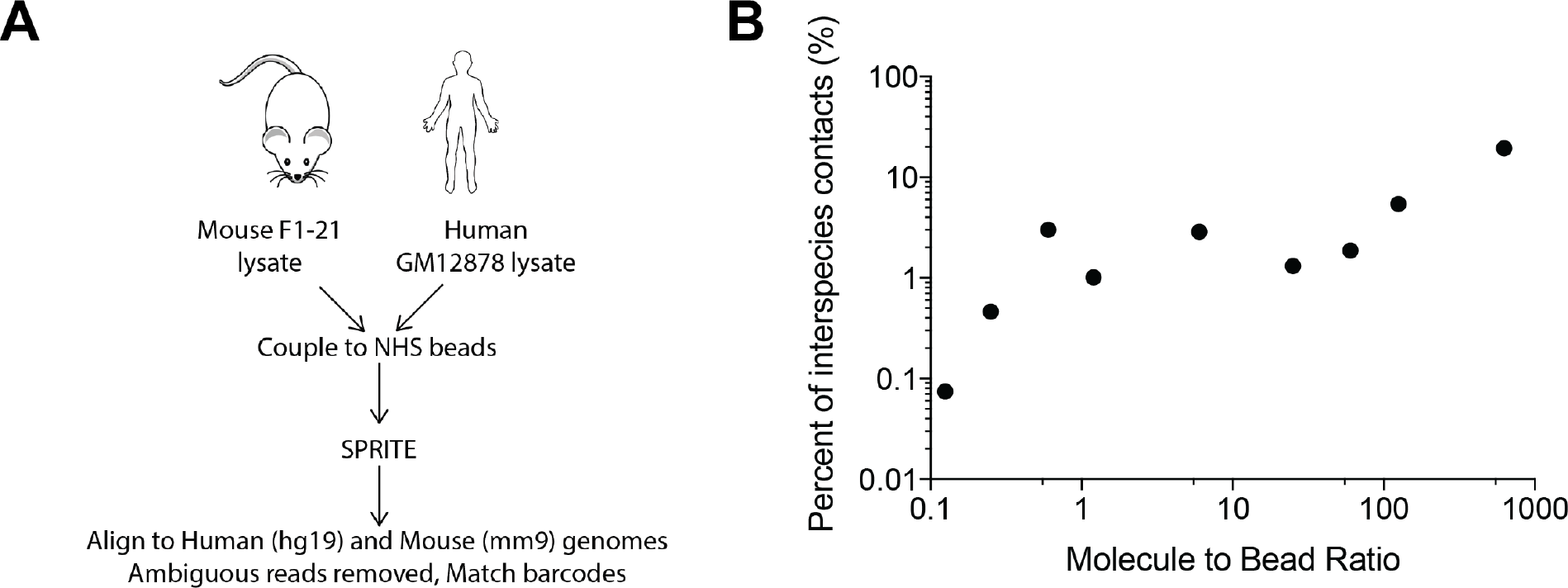
(A) Workflow of human-mouse mixing experiment to estimate noise. Lysates were separately crosslinked and lysed and combined during coupling to NHS-activated beads. After performing SPRITE, reads were aligned to both hg19 and mm9. (B) Percentage of contacts observed between human and mouse normalized by the expected number of contacts at various molecule to beads ratios.

### SPRITE tag design

All sequence tags were designed to contain at least 4 mismatches from all other tags to prevent incorrect assignments due to sequencing errors. The 5’ end of each sequence tag was designed with a modified phosphorylated base (IDT) to enable ligation. To obtain dsDNA tags, ssDNA top and bottom strands of the barcoded tags were annealed in 1x Annealing Buffer (100mM TrisHCl pH 7.5, 2M LiCl, 2mM EDgTA) by heating at 90°C for 2 minutes and slowly cooled to room temperature by reducing 1°C every 10 second in a thermocycler.

***Framework of barcoding scheme.*** In order to enable an arbitrarily large number of tags to be added to DNA, we designed a scheme that enabled reuse of the same set of barcodes. In this scheme, a universal acceptor adaptor is ligated to all DNA ends (DPM) or RNA ends (RPM). These universal adaptors contain the same sticky end overhang that is complimentary to the 5’ end of a set of adaptors referred to as “Odd” adaptors. These Odd adaptors contain a unique 3’ sticky end that is recognized exclusively by a set of “Even” adaptors, which contain a 3’ sticky end that is complementary to the Odd adaptors. In this scheme, the number of tags can be increased to as many rounds as needed, but eliminates chimera formation within a single round of split-pool tagging. We explain each of the adaptor designs in greater detail below and sequences of all tags are in **Table S5**.

***DNA Phosphate Modified (DPM) tag.*** The 5’ end of the top and bottom strands of the DPM tag molecules have a modified phosphate group that allows for ligation to dA-tailed genomic DNA and subsequent ligation of the Odd tag. DPM contains a 9-nucleotide sequence that is unique to each of the 96 DPM tags (purple region). Each DPM adaptor contains a sticky end overhang that ligates to the Odd set of adaptors (green region). The DPM sequence also contains a partial sequence that is complementary to the universal Read1 Illumina primer, which is used for library amplification (gray region).

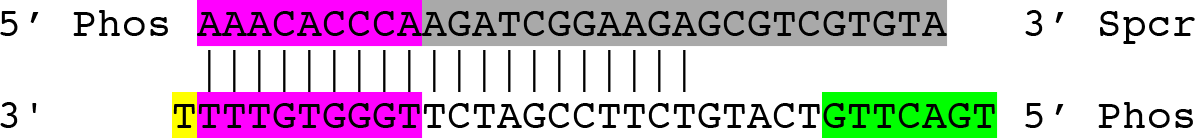

Because the DPM adaptor will ligate to both ends of the double stranded DNA molecule, we designed the DPM adaptor to ensure that we would only read the barcode sequence from one sequencing read, rather than both. To achieve this, we included a 3’ spacer on the top strand. This prevents the top strand of the Odd tag from ligating to genomic DNA. This modification is also critical for successful amplification of the barcoded DNA by preventing hairpin formation of the single stranded DNA during the initial PCR denaturation because otherwise both sides of the tagged DNA molecule would have complementary barcode sequences.

***“Odd” and “Even” Tags.*** We designed two sets of barcodes called the “Odd” and “Even” set. Both the Odd tags and Even tags have modified 5’ phosphate groups to allow for ligation. The Even adaptors are designed to have a sticky end that anneals to the Odd adaptors and the Odd adaptors is designed to contain a sticky end that anneals to the Even adaptors. The Odd adaptors are ligated in the 1^st^, 3^rd^, 5^th^,…rounds of the SPRITE process and the Even tag is ligated 2^nd^, 4^th^, 6^th^,…rounds of SPRITE. Each of the Even and Odd adaptors contain a 17nt unique barcode sequence.

***Terminal Tag.*** The terminal tag contains a sticky end that ligates to the Odd adaptors, though a terminal tag can also been designed to ligate to the Even adaptor. The terminal tag only contains a modified 5’ phosphate on the top strand. The bottom strand contains a region (grey) that contains part of the Illumina read 2 sequence, which allows for priming and incorporation of the full-length barcoded Read2 Illumina adaptor. The terminal tag contains a 9nt barcoded sequence.

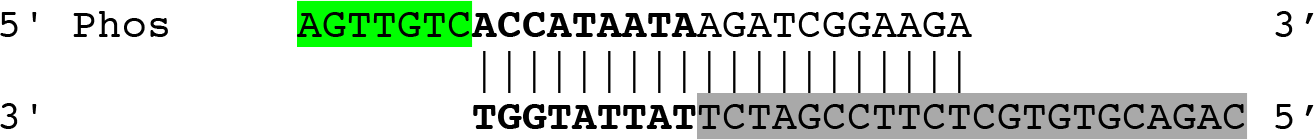

***Final DNA structure.*** After SPRITE, genomic DNA contains a DPM ligated on both ends as well as the barcodes and a terminal tag. The product is represented below:

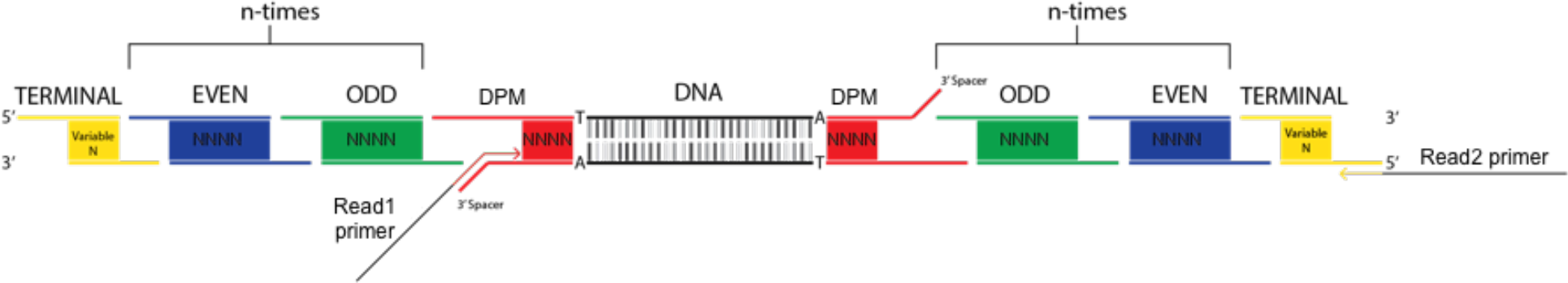

***RNA Barcoding.*** For RNA tagging, we use the same approach as above, except the first ligation to the RNA is an RNA phosphate modified (RPM) tag. The RPM tag is designed with a ssRNA overhang to specifically ligate RNA molecules using a single-stranded RNA ligase. This RNA-specific ligation tags RNA molecules to distinguish a molecule as RNA, rather than DNA, on the sequencer. The RPM adaptor contains a distinct sequence relative to DPM (pink region) and serves as a RNA-specific tag to mark each read as RNA. However, the sticky end is identical to that contained on the DPM tag (green sequence) to enable barcoding of both DNA and RNA simultaneously. The bottom strand of the RPM tag is phosphorylated after ligation of the RPM tag to RNA to ensure that the RPM tags do not form chimeras and ligate to each other during the single stranded RNA RPM ligation step. The 3’ spacer on the top strand of the RPM tag prevents ligation of single-stranded RPM molecules from forming chimeras.

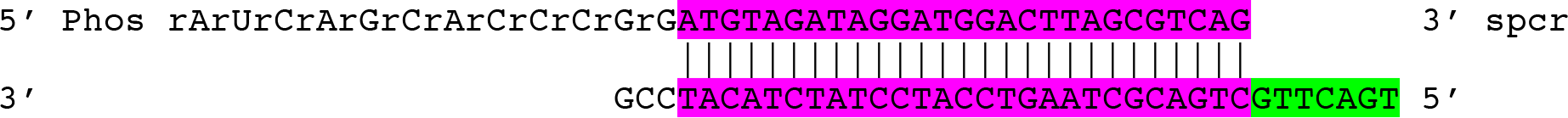

***Final RNA structure.*** The final RNA product after barcoding contains the RPM, barcodes, and terminal tag on the 3’ end of the RNA.

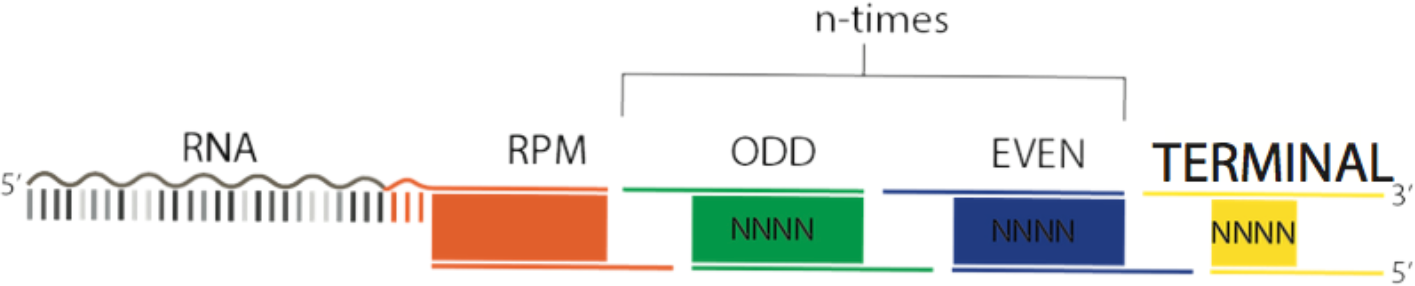

***cDNA tag.*** In order to PCR amplify cDNA, we ligate a cDNA tag to the 3’ end of all cDNA molecules. The cDNA tag contains a sequence that is part of the Illumina Read 1 primer. It is 5’phosphate modified to ligate to the 3’end of cDNA, and contains a 3’spacer to prevent chimeras of tags.

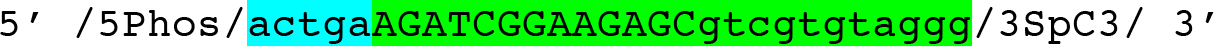

***Final Library Amplification Primers.*** DNA and RNA libraries are amplified using common primers that incorporate the full Illumina sequencing adaptors. These are Read 1 primer (AATGATACGGCGACCACCGAGATCTACACTCTTTCCCTACACGACGCTCTTCCGATCT) and Read 2 primer (CAAGCAGAAGACGGCATACGAGATGCCTAGCCGTGACTGGAGTTCAGACGTGTGCT CTTCCGATCT). The Read 1 primer amplifies the top strand of the DPM tag of DNA and cDNA tag of RNA and adds the Illumina Read 1 sequence to the molecule. The Read 2 primer amplifies the terminal tag and adds the Illumina Read 2 sequence to the molecule.

### SPRITE Data Processing and Cluster Generation

All SPRITE data was generated using Illumina paired-end sequencing on the HiSeq 2500 or NextSeq 500. Read pairs were generated with at least 115 x 100 bps. The reads have the following structure: read 1 contains genomic DNA positional information and one barcode, and read 2 has the remaining barcodes. See Table M1 for number of human and mouse reads during all steps of filtering prior to cluster generation.

***Barcode identification***. SPRITE barcodes were identified by parsing the first DNA adapter barcode sequence from the beginning of Read 1 and the remainder of the barcode sequences were parsed from Read 2. We aligned these barcode sequences against the known sets of DPM, Odd, Even, and terminal tag barcodes that were added to these samples. We allowed for up to 2 mismatches in the alignments to each internal barcode of the Odd and Even barcodes to account for possible sequencing errors. Because the barcodes were designed to contain at least 4 mismatches to any other barcode sequence, this enables for robust error correction. For DPM and terminal tag barcode alignments, we did not tolerate any mismatches due to their shorter barcode sequences. We excluded any reads that did not contain a full set of all ligated barcodes (DPM, Odd, Even, Odd,…, terminal tag) in the order expected from the experimental procedure. We also excluded all read pairs where we could not unambiguously map the barcode.

***Alignment to genome.*** We aligned each read to the appropriate reference genome (mm9 for mouse and hg19 for human) using Bowtie2 (v2.3.1) with the default parameters with the following deviations. We trimmed the 11 base pair barcode sequence (DPM) from Read 1 using the --trim5 11 parameter. To account for a short genomic fragment that might lead to additional barcode sequences being included on the 3’ end, we used a local alignment search (--local). The corresponding SAM file was sorted and converted to a BAM file using SAMtools v1.4. We stored the barcode string in the name of each record.

***Repeat Masking and Filtering Low Complexity Sequences***. We filtered the resulting BAM file for low quality reads, multimappers, and repetitive sequences. First, we removed all alignments with a MAPQ score less than 10 or 30 (heatmaps were generated for both). Second, we removed all reads that had >2 mismatches to the reference genome. Third, we removed all alignments that overlapped a region that was masked by Repeatmasker (UCSC, milliDiv < 140). Fourth, we removed any read that aligned to a non-unique region of the genome by excluding alignments mapping to regions generated by the ComputeGenomeMask program in the GATK package (readLength=35nt mask). In the human maps, all reads that overlap with an annotated *HIST* gene were removed for analysis of the histone locus in **Figure 2**.

**TABLE M1:**
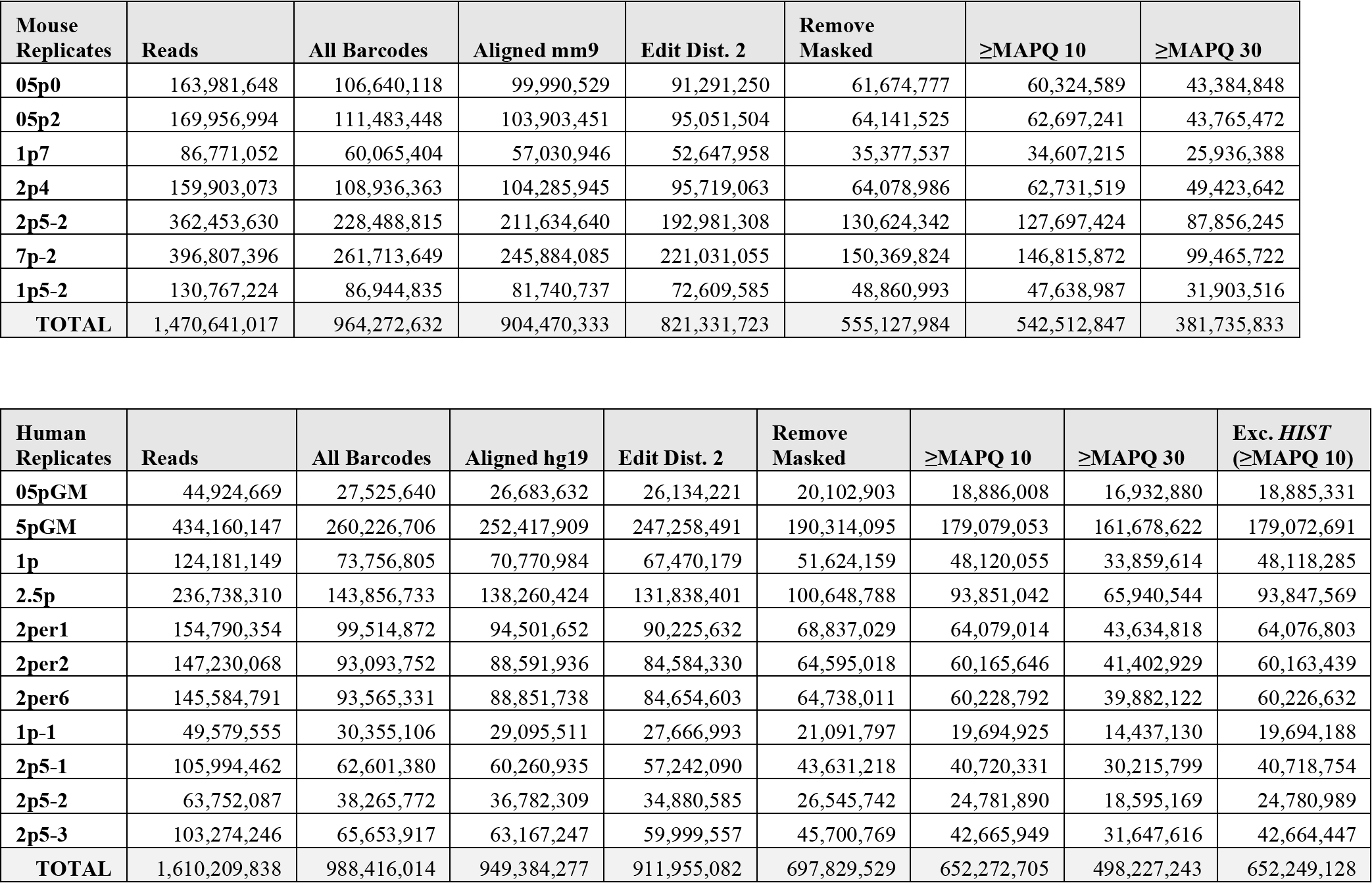
SPRITE Reads during all steps prior to cluster generation

***Defining SPRITE clusters***. To define SPRITE clusters, we grouped all read alignments that had the same barcode sequence into a single cluster. We generated a SPRITE cluster file for all subsequent analyses where each cluster occupies one line of the resulting text file containing the barcode name and genomic alignments.

***RNA-DNA SPRITE analysis***. RNA and DNA sequences were separated by the presence of a cDNA tag or DPM tag in the first 9 nucleotides of the read 1 sequence, respectively. RNA-tagged reads, identified by the cDNA tag, were aligned to ribosomal RNA sequences (28S, 18S, 5S. 5.8S, 4.5S, 45S) as well as other RNAs of interest such as snoRNAs, snRNAs including splicosomal RNAs, and Malat1. Any RNA sequences that did not align to this set of RNA genes of interest were aligned to the mm9 genome using bowtie2. DNA-tagged sequences, containing the DPM tag, were aligned to mm9. All RNA and DNA reads were subsequently filtered by edit distance and MAPQ score as described above; DNA reads were additionally filtered with a DNA mask file.

**TABLE M2:**
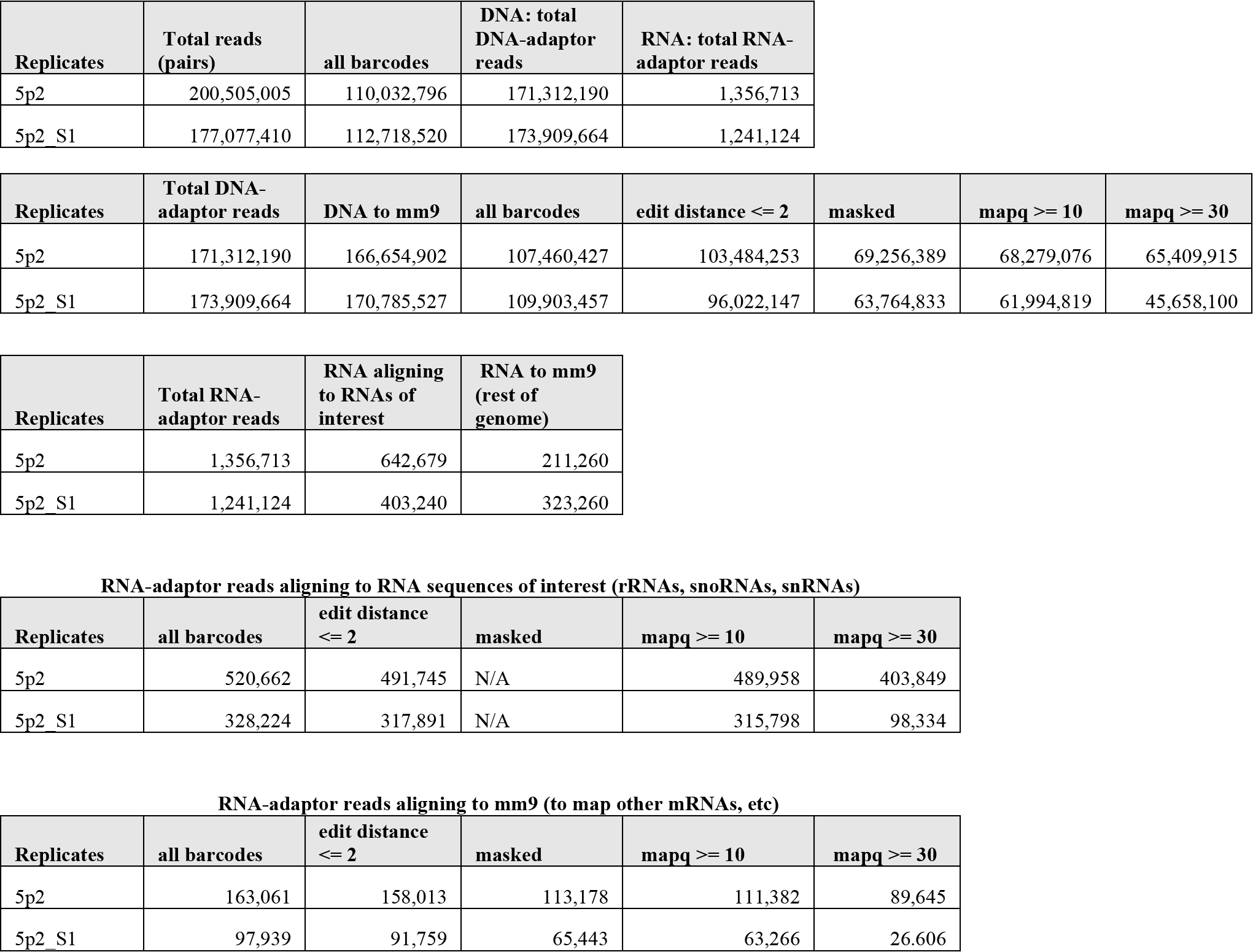
SPRITE reads in all steps prior to all steps of cluster generation for RNA-DNA tagged libraries. RNA and DNA reads, identified by a read containing an RNA-specific or DNA-specific adaptor are each processed separately.

SPRITE processing details and scripts for performing this processing are available at: www.lncRNA.caltech.edu/SPRITE/

### Generating pairwise contacts from SPRITE Data

To compute the pairwise contact frequency between genomic bins i and j, we counted the number of SPRITE clusters that contained reads overlapping both bins. We counted the number of unique SPRITE clusters overlapping the two bins and not the number of reads contained within them. In this way, if a SPRITE cluster contains multiple reads that mapped to the same genomic bin, we only count the SPRITE cluster once. Because the number of pairwise contacts scales quadratically based on the number of bins (*n*) contained within a SPRITE cluster, larger clusters will contribute a disproportionally large number of the contacts observed between any two bins. To account for this, we reasoned that a minimally connected graph containing all *n* bins would contain *n*-1 contacts. Therefore, we down-weighted each of the *n*(*n*-1)/2 contacts in a cluster by 2/*n*. In this way, the total contribution of a cluster is proportional to the minimally connected edges in the graph. This also ensures that the number of pairwise contacts contributed by a cluster is linearly proportional to the number of bins within a cluster.

Pairwise contacts were computed using multiple different bin sizes (10kb, 20kb, 25kb, 40kb, 50kb, 200kb, 250kb, 1Mb) to generate contact maps at different genomic resolutions. SPRITE contact maps were normalized by read coverage using Hi-Corrector^103^. Contacts occurring within the same bin (i.e. along the diagonal, i=j) were not considered to avoid any chance of possible PCR duplicates generating false-positive interactions.

### Comparison of SPRITE and Hi-C data

Hi-C contact maps for mouse embryonic stem cells^26^ and human GM12878 cells^23^ were normalized by read coverage using Hi-Corrector^103^.

***Compartments.*** We identified A and B compartments and insulation scores for SPRITE and Hi-C using *cworld* (Dekker lab, https://github.com/dekkerlab/cworld-dekker). To calculate compartment eigenvectors, we used the *cworld* script “matrix2compartment.pl” with default parameters with contact maps binned at 1Mb resolution as input. For human chromosomes, compartment eigenvectors were calculated separately for each chromosome arm.

***TADs.*** To calculate insulation scores, we used the *cworld* script “matrix2insulation.pl”. For human insulation scores, we used the parameters “--ss 100000 --im iqrMean --is 600000 --ids 400000” with contact maps binned at 50kb resolution. For mouse insulation scores, we used the parameters “--ss 80000 --im iqrMean --is 480000 --ids 320000” with contact maps binned at 40kb resolution.

***Loops.*** We performed aggregate peak analysis (APA) on mouse and human contact maps in both HiC and SPRITE data binned at 10kb resolution as previously described^23^. Positions for loops with end point separated by at least 200kb were obtained from mouse CH12-LX cells (1493 loops) and human GM12878 cells (5789 loops)^23^. Aggregate contact maps were computed for the median contact frequency in regions +/- 200kb of the loops.

### Analysis of higher-order *k*-mer interactions

We enumerated all higher-order *k*-mers represented in the SPRITE data at 1 megabase (Mb) resolution. We retained *k*-mers that were observed in at least 5 independent SPRITE clusters. We found that the greatest determinant of *k*-mer frequency, similar to pairwise frequency, was linear genomic distance. Accordingly, to assess the significance of a given *k*-mer, we compared the observed frequency of a given *k*-mer to the expected frequency of *k*-mers containing the same genomic distance. To do this, for a given *k-*mer, we computed the genomic distance separating each region in the *k*-mer and randomly sampled regions across the genome containing the same linear genomic distances and computed the number of SPRITE clusters containing these *k*-mers. SPRITE cluster counts were normalized by cluster size to define a weighted score and prevent large SPRITE clusters from dominating the number of *k*-mer observations. For *k-*mers of interest, enrichment was defined as the observed weighted SPRITE counts divided by the average across 100 random permutations. Genome-wide analysis was performed across 10 random permutations to identify an initial subset of enriched *k-*mers. We also retained the number of permutations that had an observed frequency larger than observed for the *k*-mer of interest and we report this percentile to rank each higher-order *k*-mer. We considered the possibility that regions with increased coverage could be observed more frequently, resulting in a high *k*-mer enrichment. However, we saw no genome-wide correlation between cumulative coverage of regions in a *k*-mer and its enrichment (R=0.19, all *k*-mers; R=0.026, enriched *k-*mers). Nonetheless, we excluded all *k*-mers containing any bins with contacts greater than 2 standard deviations from the mean.

### Comparison of SPRITE contacts in different cluster sizes

We separated SPRITE clusters into four groups: clusters with 2 to 10 reads, 11 to 100 reads, 101 to 1000 reads and clusters with 1001 or more reads. Contact maps were generated separately for each group of clusters as described above but without down-weighting for cluster size. We analyzed the relationship between genomic distance and contact frequency by computing the average contact frequency between bins separated by 40kb, 80kb, 120kb and so forth up to 100Mb. To compare heatmaps across different cluster sizes, we normalized contact frequencies by the maximum observed value such that overall contact frequency ranges from 0 to 1 for each cluster size. To examine A compartment interactions in different cluster sizes, we calculated the average contact frequency between all 1Mb bins on mouse chromosome 2 and the bins within a 9Mb A compartment region (25 to 34Mb) and normalized these values to range from 0 to 1 for each cluster size. These contact frequencies were down-weighted by cluster size (as described above).

### Analysis of inter-chromosomal interactions

To identify significant inter-chromosomal interactions, we removed all intra-chromosomal contacts. We then calculated an interaction *p*-value using a one-tailed binomial test where the expected frequency assumes a uniform distribution of inter-chromosomal contacts. We used contact maps binned at 1Mb resolution based on SPRITE clusters containing 2 to 1000 reads without down-weighting for cluster size. We built a graph where nodes represent a 1Mb bin and edges represent connections between 2 bins. We filtered edges to reduce potentially spurious contacts that may be caused by outlier bins by looking for consistency of contacts across at least 3 consecutive bins. Specifically, we only included an edge between two bins (i and j) when the edge connecting i and j was significant and all interacting pairs i±1 and j±1 were also significant. This approach produced two networks of inter-chromosomal interactions that were defined as the inactive hub (nucleolar hub) and active hub (active hub).

To identify features that distinguish these hubs and the rest of the genome, we calculated various properties, such as average gene density and average number of Pol II ChIP-seq peaks for each genomic region in these hubs and compared these to a set of control regions not contained in either hub, but with the same with the same distribution of lengths.

### Defining genomic regions near ribosomal DNA clusters

The precise location of ribosomal DNA genes in both mouse and human are unknown because they are not mapped in the reference genomes. However, approximate locations have been reported based on non-sequencing methods. In mouse, rDNA genes are encoded from the centromere-proximal regions of chromosomes 12, 15, 16, 18, and 19^73–75^. In human, rDNA genes are encoded on chromosomes 13, 14, 15, 21 and 22^33,76^. Importantly, the locations of rDNA genes can be strain-dependent in mice^73–75^,104. For instance, on chromosome 15, the *129* allele of the F1-21 hybrid mouse line does not include rDNA genes, while it does on the *Cast* allele^104^. This hybrid line was used for all DNA-DNA mapping methods and nucleolar hub identification. Instead, we performed our rRNA-DNA maps in another mouse cell line (pSM33), which is derived from a *C57BL/7* x *129SV* mouse, which are reported to contain rDNA genes on both alleles of chromosome 15. This difference in the rDNA locations between strains may explain why we observe a stronger enrichment of rRNA on chromosome 15 in the rRNA-DNA maps than in the DNA nucleolar hub contact maps from the F1-21 hybrid mice (Figure 4A).

### RNA localization quantification

To quantify the localization of ribosomal RNA (rRNA) across the genome, we split SPRITE clusters into two groups, one with clusters that contained at least 1 rRNA read (rRNA positive clusters) and the other with clusters lacking any rRNA reads (rRNA negative clusters). We then calculated the ratio of rRNA positive clusters to rRNA negative clusters for each 1Mb bin, normalizing for total number of clusters in each group, and defined this ratio as the rRNA enrichment for each bin. We calculated rRNA enrichment *p*-values for each bin using a one-tailed binomial test where the expected frequency was based on the rRNA negative clusters. To quantify Malat1 and U1 localization, we obtained Malat1 and U1 RAP-DNA alignments from Engreitz *et al*.^79^ and calculated Malat1 and U1 enrichment for each 1Mb bin by normalizing to input RAP-DNA alignments.

### DNA-FISH

DNA fluorescence in situ hybridization (DNA-FISH) was performed with Agilent SureFISH DNA-FISH probes following the manufacturer’s protocol with adaptations noted below. Female F1-21 Cells were fixed on coverslips in a 24 well plate using 300uL of Histochoice for 10 minutes at room temperature, then dehydrated through incubation in a series of graded ethanol concentrations up to 100% ethanol, and air-dried. The coverslips were then mounted on a 5uL mixture of probe set targeting a selected DNA locus (Agilent) and SureFISH Hybridization Buffer (Agilent). The coverslips and probe mixture were denatured for 8 minutes at 83°C, then incubated at 37°C overnight. The following morning coverslips were washed with FISH Wash Buffer 1 (Agilent) at 73°C for 2 min on a shaking incubator at 300rpm, and FISH Wash Buffer 2 (Agilent) at room temperature for 1 minute. Coverslips were then rehydrated and suspended in 1X PBS in preparation for immunofluorescence staining. Probe sets used for FISH were designed by Agilent technologies using their standard procedures against the following genomic regions (Table M3).

**TABLE M3:**
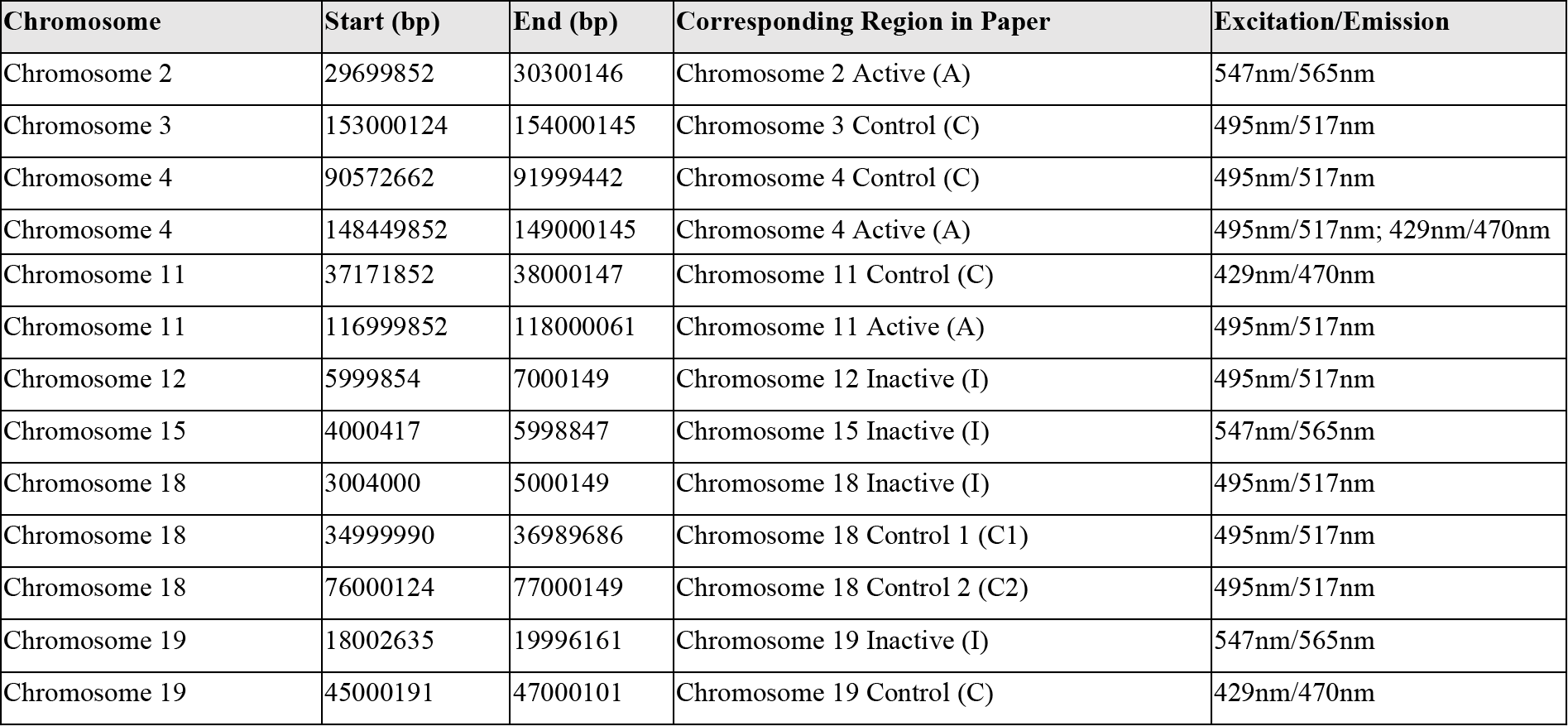
DNA FISH probes used in this paper.

### Immunofluorescence

Coverslips were permeabilized with 0.1% Triton-X in PBS at room temperature for 10 minutes, then blocked with 1X blocking buffer (Abcam) in PBS at room temperature for one hour. The coverslips were then incubated with primary antibodies in a humidified chamber at room temperature for one hour. The coverslips were washed with 0.1% Triton-X in PBS at room temperature, then incubated with secondary antibodies in a humidified chamber at room temperature for one hour. Coverslips were washed with PBS and H_2_O and mounted on slides in ProLong^®^ Gold antifade reagent with DAPI (Life Technologies). The primary antibodies used for IF were rabbit polyclonal anti-Nucleolin (Abcam; ab22758; 1:1000) and mouse monoclonal anti-SC35 (Abcam; ab11826; 1:200). The secondary antibodies used for IF were Alexa Fluor® 647 goat anti-rabbit IgG (H+L) (Life technologies; A21244; 1:300) and DyLight® 650 goat anti-mouse IgG (H+L) (Bethyl; A90-116D5; 1:300).

### Microscopic imaging

DNA FISH/IF samples were imaged using a Leica DMI 6000 Deconvolution Microscope, with a z-stack collected for each channel (4 μm; step size, 0.2 μm). The objectives used were the Leica HC PL APO 63x/1.30 GLYC CORR CS2 objective and the Leica HCX PL APO 100X/1.40-0.70na OIL objective. Samples were also imaged with a ZEISS Laser Scanning Microscope (LSM) 800 with the ZEISS i Plan-Apochromat 63x/1.4 Oil DIC M27 objective, with a z-stack collected for each channel (step size, 0.37 μm). Deconvolution was performed using Huygens Professional version 17.04 (Scientific Volume Imaging, The Netherlands, software available at http://svi.nl) using the built-in theoretical point spread function, the classic maximum likelihood estimation (CMLE) algorithm, a signal to noise ratio of 20, and 50 iterations.

### Calculating distance between DNA loci

The nuclei of individual cells were identified by DAPI staining, and cells containing two spots per DNA-FISH channel were identified manually. Images were cropped to only contain the identified cell. Analysis of cells in three dimensions was performed using Imaris version 8.4.1 (Bitplane Inc, software available at http://bitplane.com) with the ImarisXT module. Both alleles for each DNA locus were defined by applying the Imaris “Spot” function (diameter = 0.5μm, background subtraction) on the corresponding fluorescent channel. The distance between DNA loci was calculated by running the XTension “Distances Spots to Surfaces” function and manually recording the smallest distance between alleles of differing loci.

### Calculating distance between DNA loci and nuclear bodies

The nucleolus and nuclear speckle, identified by immunofluorescence of nucleolin and SC35 respectively, were defined in Imaris by performing the Imaris “Surface” function (detail = 0.126 μm, absolute intensity). Custom Imaris XTensions were used to calculate a distance transform approximating Euclidean distance for the region outside of the generated surface (“Batch Process Function” by Pierre Pouchin and “Distance Transformation Outside Object For Batch” by Matthew Gastinger, obtained on open.bitplane.com). The edges of the surface served as boundary voxels; regions inside the surface were assigned a distance transform value of 0. The distance of an allele to the nucleolus or nuclear speckle was defined as the minimum distance transform value of the corresponding spot, from the edge of the surface to the nearest edge of the DNA-FISH sphere.

### Comparison of SPRITE and DNA FISH

To compare SPRITE and DNA FISH measurements, we used SPRITE contact frequencies from contact maps binned at 1Mb resolution based on SPRITE clusters containing 2 to 1000 reads without down-weighting for cluster size. We note that similarly high correlations between SPRITE and DNA FISH measurements were observed when large SPRITE clusters (>100 reads) are included in the analyses (consistent with the observation of inter-chromosomal interactions in larger clusters in Figure 3C), both with and without weighting for cluster size (Table M4). SPRITE contact frequencies were obtained for 1Mb bins that overlapped with each DNA FISH probe region and compared to DNA FISH distance measurements with the corresponding probe region.

**TABLE M4:**
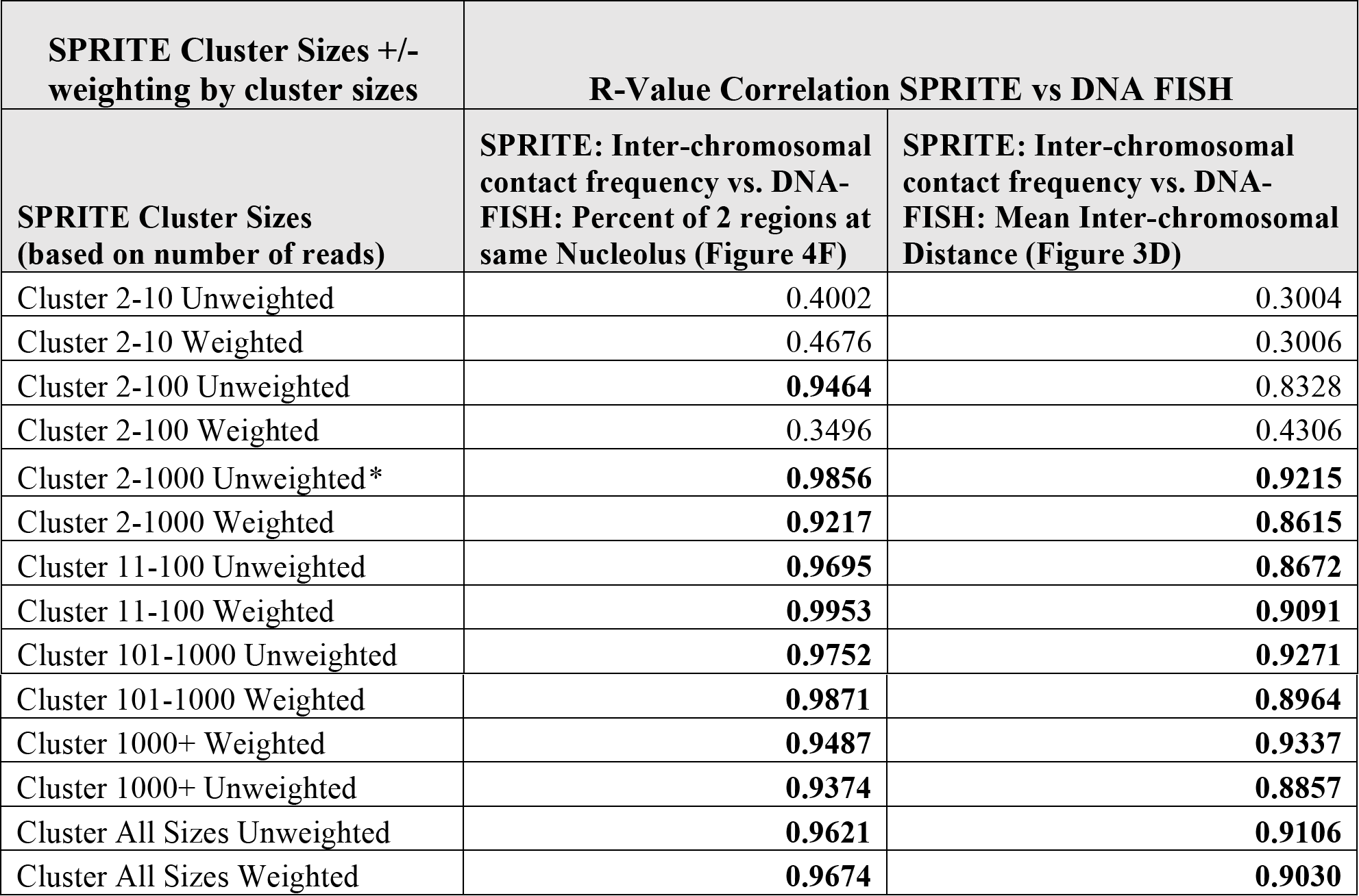
SPRITE highly correlated with DNA FISH for most cluster sizes, regardless of weighting or no weighting by cluster size. R-values >0.85 in bold. * represents the values currently shown in the analyses for Figures 3D and 4F.

### SPRITE contact frequency with the nucleolar and active hubs

We defined contact frequency with the nucleolar and active hubs for each 1Mb bin based on the average inter-chromosomal contact frequency with all regions in the nucleolar and active hub, respectively. P-values were calculated for these contact frequencies using a one-tailed binomial test where the expected frequency assumed a uniform distribution of inter-chromosomal contacts. Contact frequencies with these hubs were compared to RNA localization (described above) and data from ChIP-seq (e.g., Pol II, H3K4me3) and GRO-seq experiments obtained from ENCODE and from Jonkers et al., 2014^91^, respectively.

